# A Compact and Effective Photon-Resolved Image Scanning Microscope

**DOI:** 10.1101/2023.07.28.549477

**Authors:** Giorgio Tortarolo, Alessandro Zunino, Simonluca Piazza, Mattia Donato, Sabrina Zappone, Agnieszka Pierzyńska-Mach, Marco Castello, Giuseppe Vicidomini

**Affiliations:** Molecular Microscopy and Spectroscopy, Istituto Italiano di Tecnologia, Via Enrico Melen, 83, Genoa, 16152, Italy; Nanoscopy and NIC@IIT, Istituto Italiano di Tecnologia, Via Enrico Melen 83, Genoa, 16152, Italy; Dipartimento di Informatica, Bioingegneria, Robotica e Ingegneria dei Sistemi, University of Genoa, Via Dodecaneso 35, Genoa, 16146, Italy; Genoa Instrument, Via Enrico Melen 83, Genoa, 16152, Italy

**Keywords:** Fluorescence Lifetime, Image Scanning Microscopy, Digital Frequency Domain, Single Photon

## Abstract

Fluorescence confocal laser-scanning microscopy (LSM) is one of the most popular tools for life science research. This popularity is expected to grow thanks to single-photon array detectors tailored for LSM. These detectors offer unique single-photon spatiotemporal information, opening new perspectives for gentle and quantitative super-resolution imaging. However, a flawless recording of this information poses significant challenges for the microscope data-acquisition system. Here, we present a data-acquisition module based on the digital frequency domain principle, able to record photons’ essential spatial and temporal features. We use this module to extend the capabilities of established imaging techniques based on single-photon avalanche diode (SPAD) array detectors, such as fluorescence lifetime image scanning microscopy. Furthermore, we use the module to introduce a robust multi-species approach encoding the fluorophore’s excitation spectra in the time domain. Lastly, we combine time-resolved stimulated emission depletion microscopy with image scanning microscopy, boosting spatial resolution. Our results demonstrate how a conventional fluorescence laser scanning microscope can transform into a simple, information-rich, super-resolved imaging system with the simple addition of a SPAD array detector with a tailored data acquisition system. We expected a blooming of advanced single-photon imaging techniques which effectively harness all the sample information encoded in each photon.

## 1 Introduction

Asynchronous read-out single-photon avalanche diode (SPAD) array detectors [1–3] are expected to terrifically extend the abilities of fluorescence laser-scanning microscopy (LSM). In contrast with the typical LSM single-element detectors, but similar to scientific cameras, this class of sensors preserves the spatial distribution of the impinging fluorescence photons. Differently from a scientific camera, every element of the array is a fully independent SPAD [4], which provides a high temporal precision (*<* 100 ps) digital signal for each detected single-photon. The combination of tailored data-acquisition (DAQ) systems with these SPAD array detectors enables photon-resolved measurement of the fluorescence emission: every photon can be tagged with (i) spatial signatures – namely, the emission and detection coordinates of the photon in the detector and sample plane, respectively – and (ii) temporal signatures – namely, the time delay of the photon with respects to specific reference events. A series of recent publications [5–10] have demonstrated the benefits of leveraging the unique information provided by the SPAD arrays. The photon-resolved spatial and temporal information improves the performances of most advanced LSM-based techniques and paves the way for new ones. Such unprecedented results laid the foundation of a new microscopy paradigm that we named *single-photon laser-scanning microscopy* (SP-LSM) [9]. The features of a particular technique based on the SP-LSM paradigm depend on the characteristics of the SPAD array detectors [11, 12] and those of the DAQ systems. The spatial information alone, provided by the detector array, has already boosted the performances of numerous techniques. Among them, confocal LSM evolved into image-scanning microscopy (ISM), capable of generating super-resolved images at an exceptional signal-to-noise ratio (SNR) level [6, 13–15]. Furthermore, detector arrays also boosted the optical sectioning capabilities of ISM [10]. Importantly, the same benefits have been easily transported to two-photon-excitation (TPE) [16] and stimulated-emission depletion (STED) microscopy [10]. Fluorescence-fluctuation spectroscopy [17] (FFS) is another technique that benefited from structured detection [3, 8], gaining access to molecular mobility information previously hidden. The aforementioned techniques require access to temporal scales of the order of the pixel dwell time or (histogram) bin time width for imaging or FFS, respectively. These experimental parameters are typically in the range of a few microseconds. Indeed, the relative DAQ systems only need to count photons – on every channel of the SPAD array – detected within each pixel of the image or bin of the histogram. However, the power of the SP-LSM paradigm is fully unleashed with the techniques requiring access to the nano or picosecond scale. In fluorescence lifetime ISM (FLISM) [6, 9], the photon-arrival time with respect to the fluorophore excitation event enables mapping the fluorescence lifetime distribution in the sample at super-resolution. Such information helps understand the properties of the bio-molecular environment [18], deciphering the bio-molecule structural changes [19], and implementing multi-species imaging [20]. In quantum ISM (Q-ISM) [5], the photon-coincidences allow the construction of an image with a spatial resolution beyond the limits of conventional ISM. Super-resolution optical fluctuation ISM (SOFISM) [7] exploits fluorescence fluctuations on the (sub-)microsecond scale to improve spatial resolution. In these cases, the DAQ system must include a multi-channel time-tagger (TT) module. This latter registers each photon detection event with spatial and temporal coordinates, namely the position on the detector plane and the detection times – absolute and relative to a series of reference signals (e.g., the sync of the laser pulses and the sync from the beam scanning system). Time-tagger DAQ (TT-DAQ) systems are the greatest ally for SP-LSM, fully preserving the single-photon information collected by the SPAD array detectors. However, they are more complex and expensive than photon-counting DAQ systems. Furthermore, they require to transfer and store a large amount of data. Time-tagger DAQ typically records signals with tens of picoseconds precision with a range of up to several seconds. While the complexity and cost can be reduced using DAQ boards based on field-programmable-gate-array (FPGA) [9, 21], the storage problem cannot be solved without sacrificing some information. A possible solution would be to design an application-specific TT-DAQ system, where the temporal characteristics and information content are bounded by the needs of the measurements, as recently done for Q-ISM [22].

In this work, we propose a valuable alternative to high-precision time-tagging DAQ systems to implement FLISM. We leverage the digital-frequency-domain (DFD) principle [23, 24] to implement an FPGA-based multi-channel DAQ system able to terrifically reduce the data transfer and storage without compromising the performance of FLISM. Specifically, we use a heterodyne scheme to estimate the photon-arrival time with a sampling period down to 400 ps. Instead of transferring the arrival time of each detection event to the computer, we build the fluorescence decay histogram directly on the FPGA board. Thus, we only need to transfer and store a histogram for each channel and imaging pixel (or histogram time bin for FFS). This strategy also allows to visualise in real-time the lifetime measurements and to easily embed on the same FPGA board both the time-resolving DAQ and the microscope control system.

We integrate our new DFD-DAQ and control system into a SP-LSM architecture equipped with a commercial SPAD array detector. We validate the DFD module embedded in the microscope control and DAQ unit by implementing different FLISM-based imaging techniques. We combine our new platform with the fluorescence lifetime phasor analysis to implement functional super-resolution imaging. Despite the lower temporal precision than a typical TT-DAQ module, the DFD-DAQ module allows the investigation of time-scales short enough to monitor the lifetime changes of molecules with biological interest. The combination of our architecture with the phasor analysis also enables super-resolution imaging of multiple fluorophores without spectral emission separation. In this case, we distinguish the different dyes on a sample by leveraging their different fluorescence lifetime. Furthermore, we took advantage of the integration of the DFD module into the microscope control system to implement a pulse-interleaving multi-wavelength excitation scheme. Different wavelengths enable to excite multiple fluorophores, which might have overlapping absorption spectra. We introduce a new phasor-based method for multi-species separation, also in the event of spectral excitation cross-talk. Finally, we implement nanoscopy by combining STED-ISM with separation-by-lifetime tuning (SPLIT) [25, 26]. This approach uses the phasor-analysis and the fluorescence lifetime changes induced by the stimulation emission process [27] to improve the resolution of STED microscopy. Our results demonstrate the versatility and vast potentialities of the proposed architecture.

## 2 Results

In this work, we used a custom fluorescence laser-scanning microscope (Fig. 1a) equipped with a 5 × 5 SPAD array detector, a pair of triggerable picosecond pulsed diode lasers for excitation (560 nm and 640 nm), and a three-dimensional scanning system. A sub-nanosecond pulsed fiber laser (775 nm), synchronised with the excitation lasers, enables stimulated emission for STED microscopy. The FPGA board includes both the multi-channel DFD-DAQ and the microscope control modules. The first records the inputs from each channel of the SPAD array detector; the latter provides the analog outputs for driving the scanner and the digital outputs for triggering the pulsed lasers. Since both modules have common clocks, their synchronisation is straightforward: the lifetime histogram is calculated on the FPGA board and returned at the end of each pixel dwell time. Furthermore, the trigger signals for the lasers can be delayed to enable pulse-interleaving excitation – namely, alternating excitation at different wavelengths. For each scan point and each channel of the SPAD array, the FPGA builds the photon-arrival histogram using 25 low-precision time-to-digital converters (TDCs) based on the DFD principle. The DFD system estimates the photon-arrival time by measuring the delay between the excitation event and the digital single-photon signal coming from the SPAD.

**Fig. 1.**
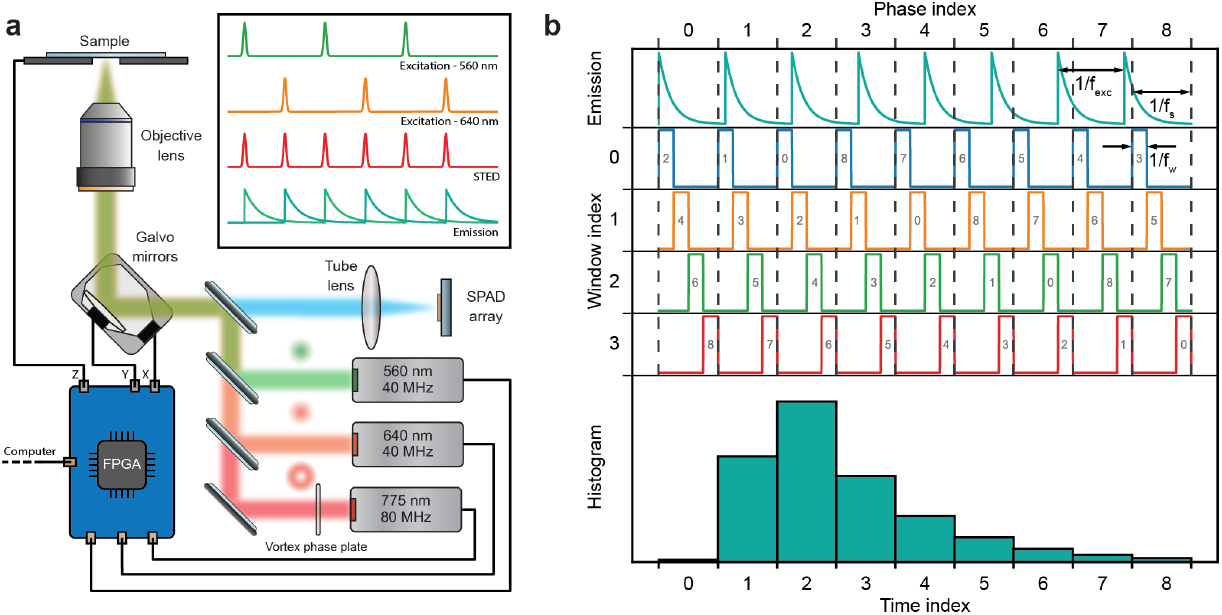
Setup and acquisition scheme. **a**, sketch of the STED-ISM custom microscope. The FPGA board controls two excitation lasers, a depletion laser, a couple of galvanometric mirrors, and a piezoelectric-controlled stage. The same FPGA board reads the output of each channel of the SPAD array detector. **b**, working principle of the heterodyne acquisition. The lifetime decay is sampled along multiple excitation periods using a detuned sampling frequency. The sampling accumulates a delay in each period, enabling data collection at different time points. Each sampling period is sub-sampled by short windows, boosting the temporal resolution.

The DFD-TDC performs digital heterodyning: excitation and sampling frequencies, *f*_*exc*_ and *f*_*s*_, are slightly different. The periodicity of the beating signal *f*_*c*_ = *f*_*s*_ *−f*_*exc*_ determines the duration of a DFD cycle. This latter spans multiple excitation cycles, time-tagging photons at different delays from the excitation pulse. More in detail, each sampling period is divided into an integer number of short windows at frequency *f*_*w*_ = 1*/T*_*w*_. Thanks to a pair of counters – the phase and window indices – the photon arrival times are remapped into a single excitation period, generating a densely sampled histogram of the fluorescence decay with a temporal resolution beyond that of conventional sampling (Fig. 1b).

We designed our system to generate the excitation and sampling frequencies from the same FPGA clock: the excitation signal triggers the laser pulses, and the temporal histogram is fully calculated at the FPGA level. In short, our system embeds the control of the microscope and the measurement of the photon-arrival time histogram in a single compact device. Advantageously, our architecture uses only a limited amount of hardware and software resources (see Supplementary Note A).

In this work, we use two specific configurations, running at *f*_*exc*_ ≈ 20 MHz and *f*_*exc*_ ≈ 40 MHz. Both implementations achieve a temporal resolution – namely, the temporal width of the histogram bin – of about 400 ps. Instead, the temporal precision is dictated by the value of *T*_*w*_, which is about 2 ns.

### 2.1 Fluorescence Lifetime Assay

To validate the proposed architecture, we used the DFD configuration running at *f*_*exc*_ ≈ 40 MHz to measure the photon-arrival time histogram *f* (*t*) (Fig. 2a top) from a sample of diluted solution of Alexa 594 with a well-known fluorescence lifetime (*τ* = 3.97 ns) [28]. Given the photon-arrival time histogram, we can estimate the fluorescence lifetime value by fitting the decay to an exponential function convolved with the system impulse-response-function (IRF). Nonetheless, the phasor approach enables a fit-free lifetime value estimation, inherently considers multi-exponential decays, and grants access to a straightforward graphical representation of lifetime values [29]. Thus, we use the photon-arrival time histogram *f* (*t*) to calculate the intensity normalised phasor vector components for the first harmonic (*g, s*) and the corresponding phase and modulation values (*ϕ, m*). We then represent the phasor vector for the Alexa 594 into a two-dimensional histogram called the phasor plot (Fig 2a top, inset). The theoretical model predicts that the phasor of single-exponential decay lies on a semicircle in the complex plane, known as the *universal circle* [30]. However, the system’s IRF alters the experimentally measured decay, whose phasor no longer lies on the universal circle. Starting from the reference lifetime value of the Alexa 594, we calculated the corrected phasor. The ratio of the measured to the correct phasor leads to the phasor components of the system’s IRF, which can be used to calibrate all successive measurements (see Supplementary Note B). Indeed, we measured a solution of Rhodamine 101, whose raw and corrected phasors are shown (Fig. 2 a). The fluorescence lifetime calculated from the phase of the calibrated phasor phase value is *τ*_*ϕ*_ = 4.16 ns, similar to the values reported in literature [31]. In our DFD implementation, the phase difference between the excitation and the sampling signal is not fixed and may vary from measurement to measurement. Thus, we need to measure the phase of an additional reference. For this, we record the trigger signal used to synchronise the excitation laser as an additional virtual channel. The calibration procedure is applied independently to each element of the SPAD array detector (Suppl. Fig. D2).

**Fig. 2.**
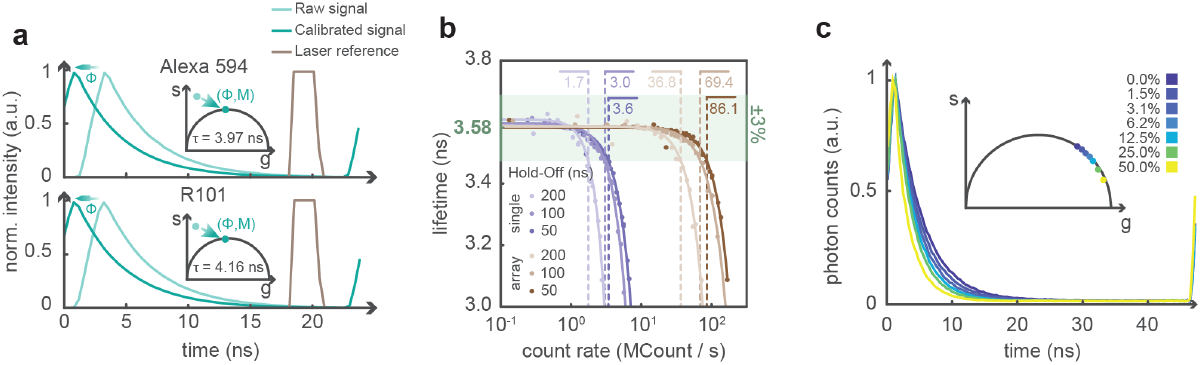
Validation of the DFD architecture. **a**, calibration procedure. The measured phasor is shifted and rescaled by the IRF calibration values. Additionally, each histogram is phase-shifted by the value provided by the trigger reference signal. Top: calibration of Alexa 594. Bottom: calibration of Rhodamine 101. **b**, Estimated lifetime of an autofluorescent plastic slide as a function of the detected photon flux and of the hold-off time of the SPAD. The reported values are the saturation thresholds (in units of MCount/s), defined as the flux at which lifetime estimation deteriorates by a factor larger than 3%. **c**, Lifetime estimation of a fluorescein solution at increasing concentration a quencher, potassium iodide. We depict with the same colors the exponential decays and the corresponding phasors.

Implementing the TDC using a DFD approach allows for recording virtually all photon signals. Indeed, the dead-time of the DFD module is given by the window duration, in our implementation *T*_*w*_ = 2 ns. Within each window, the DAQ system can only discriminate between the presence or absence of the detection event. Nonetheless, state-of-the-art SPAD array detectors have hold-off times in the range of tens of nanoseconds. Thus, the dead time of the DFD module does not introduce any additional practical limitation. However, the low speed of the USB 2.0 interface, which connects the card to the computer, constraints the data-transfer rate of our implementation of the multi-channel DFD-DAQ system. Assuming a depth of 16 bits for each time-bin of the histogram and 26 parallel channels, USB 2.0 is able to transfer the whole histogram every 125 μs, imposing a lower limit on the pixel dwell time.

We experimentally verified the maximum photon flux that can be sustained by our multi-channel DFD-DAQ module. We registered the fluorescence signal generated by an autofluorescent plastic slide (Chroma) at increasing excitation power. We calculated the fluorescence lifetime *τ*_*ϕ*_ from the histograms retrieved at increasing photon-fluxes (Fig. 2b). The fluorescence lifetime of the slide does not depend on the excitation power. Thus, a reduction of the estimated fluorescence lifetime value indicates the saturation of the DAQ system. Indeed, at high count rates, the early photons are most likely to be detected because of the hold-off of the detector. This phenomenon, known as the pile-up effect, causes the appearance of an additional short-lived exponential component and a decrease of the measured lifetime. In these experiments, the photons are not distributed uniformly across the whole detector array, but according to the diffraction laws. Specifically, light spreads according to the fingerprint distribution [10], namely a bell-shaped function peaked in the central element of the detector array. The central element starts saturating (namely, the lifetime error is greater than 3%) above 1.7 megacounts/s. To measure the saturation of the whole system, we integrated all photon-arrival histograms before calculating the fluorescence lifetime. We find the saturation threshold of the whole system to be at 36.8 megacounts/s. The dead time of the SPAD elements is the major cause of saturation. Thus, reducing the hold-off time of the SPAD array from 200 ns to 50 ns, the saturation threshold shifts toward higher photon-fluxes: 3.6 megacounts/s for the central element and 86.1 megacounts/s for the whole system.

Finally, we used our platform to measure the fluorescence lifetime values for a series of fluorescein solutions obtained with increasing concentrations of the quencher potassium iodide. As expected, the platform retrieves shorter lifetimes as the quencher concentration increases (Fig. 2c).

### 2.2 FLISM for Functional Imaging

After validating the multi-channel DFD-DAQ module, we used the module to implement fluorescence lifetime image scanning microscopy (FLISM) [6, 9]. FLISM harnesses the photon-resolved spatiotemporal information provided by the SPAD array to obtain a super-resolved fluorescence lifetime map of the sample. On each acquisition, the microscope builds a 5D photon-counts map *i*(**x**_*s*_, **x**_*d*_, *t*). We reconstruct a super-resolution image from the raw data using adaptive pixel reassignment (APR) as in conventional ISM. In FLISM, we iterate the APR analysis on each temporal bin, obtaining a time-resolved super-resolution image *i*_FLISM_(**x**_*s*_, *t*). We apply phasor analysis to this latter to obtain the fluorescence lifetime map *τ* (**x**_*s*_). We started by imaging fixed HeLa cells in which we stained Tubulin with ATTO647N (Fig. 3a) and nuclear-pore complexes with Abberior STAR 580 (Fig. 3b). We used the excitation wavelength and detection filter that best matches with the probes. For both structures FLISM outperforms conventional confocal laser scanning microscopy (CLSM). The FLISM images show a better resolution than the open-pinhole confocal microscopy counterpart (Suppl. Fig. D3 and Suppl. Fig. D4). When closing the pinhole, the optical resolution of confocal microscopy should reach the FLISM resolution. However, the pinhole rejects most of the fluorescence photons, drastically reducing the image’s signal-to-noise ratio (SNR). The SNR is a crucial parameter for fluorescence lifetime imaging. Indeed, regardless of the method used to analyse the histogram arrival time, the worst is the SNR, and the worst is the precision of the fluorescence lifetime estimation. The differences in fluorescence lifetime precision between the three imaging modalities (open-pinhole, closed-pinhole and ISM) are evident when comparing the respective fluorescence lifetime maps and the phasor plots (Fig. 3a,b). Since the tubulin network and the nuclear pore complexes are stained in fixed cells, the fluorescence lifetime should be constant across the whole sample. A potential source of lifetime heterogeneity in the sample could be the high concentration of fluorophores: in this condition, fluorophores can self-quench each other, resulting in shorter lifetime values [27]. Here, we assume that self-quenching and similar phenomena are negligible. Thus, any variation of the fluorescence lifetime values between the different pixels is induced by an estimation error. Closed-pinhole images show different pixels with out-of-range fluorescence lifetime values, indicating a low precision. Comparing the fluorescence lifetime histograms and the phasor plots confirms the same trend: closed-pinhole confocal microscopy shows a broader distribution than open-pinhole confocal microscopy and ISM (Suppl. Fig. D5). The effect, known as super-brightness [32], makes the ISM image’s SNR even better than that of open-pinhole confocal microscopy, further enhancing the precision of FLISM.

**Fig. 3.**
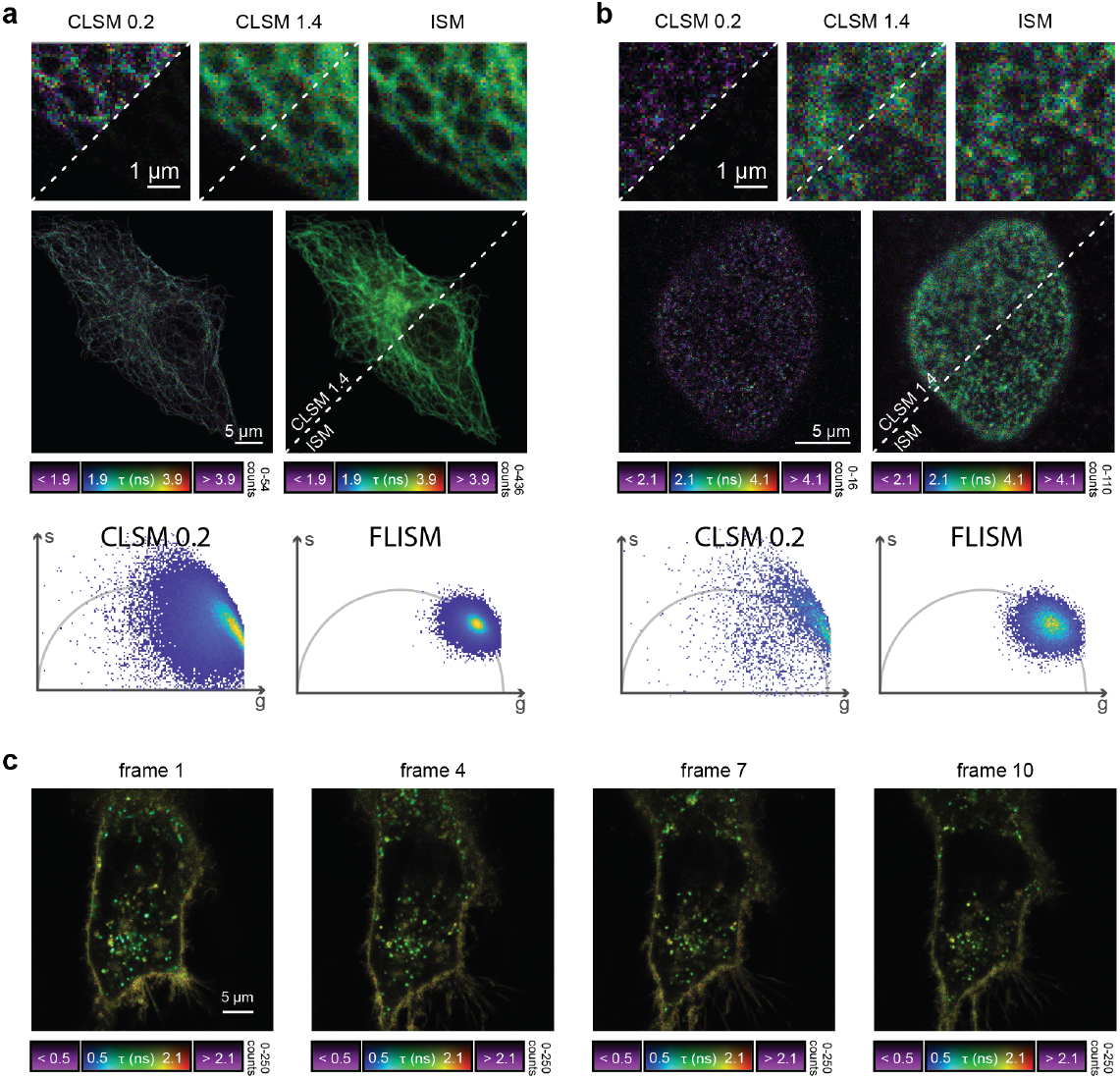
FLISM imaging. **a**, Alpha-tubulin stained with ATTO647N in fixed HeLa cells. **b**, nuclear pore complexes stained with Abberior STAR 580 in fixed HeLa cells. **c**, living HeLa cell labeled with the CellBrite®NIR680 cytoplasmic membrane dye. The displayed frames are captured at *t* = 0 min, *t* = 7.5 min, *t* = 15 min, *t* = 22.5 min.

Next, we used the FLISM approach to measure living HeLa cells labelled with the CellBrite® NIR 680 cytoplasmic membrane dye. Because the dye uniformly labels any membrane structure, conventional intensity-based imaging cannot distinguish the signal stemming from cell membrane or membrane-enclosed organelles (e.g., lysosomes). Conversely, FLISM reveals a broad distribution of the fluorescence lifetime values, which allows for distinguishing the plasma membranes (longer fluorescence lifetime) from intracellular vesicles or lipid-based structures (shorter fluorescence life-time). Indeed, the variation of the probe’s fluorescence lifetime allows for monitoring the changes in the local environment. The superior SNR of ISM imaging allows for working at reduced illumination intensity, thus allows for long-term imaging (Fig. 2c). We follow the variations of the fluorescence lifetime of the membrane dye within the HeLa cells for more than two hours.

### 2.3 FLISM for Multi-Species Imaging

Biological investigations often require visualising multiple bio-molecules and cellular compartments simultaneously. Multi-species imaging generally targets the different bio-molecules with spectrally separable fluorophores. A time-resolved detection system enables separating fluorophores based on their fluorescence lifetime, offering a viable alternative to distinguish dyes with similar emission and excitation spectra. Conveniently, this solution requires only a single detector. Furthermore, lifetime separation does not preclude spectral separation. Indeed, the two approaches can be combined to enable the separation of an even wider number of species [20].

Here, we demonstrate how our DFD-DAQ module can implement imaging through fluorescence lifetime multiplexing. In particular, we show that FLISM method can perform super-resolution multi-species imaging using a single SPAD array detector. We stained *α*-tubulin and lamin A in fixed MCF10A cells with Abberior STAR RED and Abberior STAR 635, respectively. The two dyes show almost identical excitation spectra and largely overlapping emission spectra. We excited both dyes with the same pulsed laser (*λ*_*exc*_ = 640 nm) and detected their fluorescence signals within the same spectral window and SPAD array detector. However, the two probes emit fluorescence with substantially different lifetimes. A straightforward approach for fluorescence life-time multiplexing imaging is phasor segmentation. Typically, two (or more) regions on the phasor plot are identified using the single-probe samples. The same regions are then reported in the phasor plot of the multi-probes image, and the point on these regions are back-projected to generate the single-species images. While easy to use, fast, and visually intuitive, the phasor-segmentation approach assumes that the same pixel contains a signal stemming only from a specific probe. However, this assumption is too restrictive in many practical scenarios. For this reason, we analyzed FLISM data with an algorithm inspired by the method known as separation-by-lifetime-tuning (SPLIT) [25]. To enable multi-species imaging, we developed a similar algorithm that decomposes the FLISM signal in each scan point **x**_*s*_ into the components associated with the two species. This method – named phasor separation – exploits the phasor representation of fluorescence decays. We describe each decay with the first frequency component of the corresponding Fourier transform – namely, the phasor. With such an approach, the mixed signal is conveniently represented as the linear combination of two phasors, one for each fluorophore. Thus, we transform a non-linear regression algorithm – such as a model-based fitting – into a linear problem that admits an easy-to-compute solution (see Supplementary Note C). We tested our phasor-separation method on the FLISM dataset of MCF10 cells. First, we apply the phasor analysis to the images of two separate samples containing only one or the other dye. Thus, we retrieve the corresponding phasor coordinates, which map to different points of the universal circle. Then, we compared the well-known phasor-segmentation and our new phasor-separation method. Both methods are feasible, but phasor segmentation results in an excessively sharp classification that does not admit the presence of multiple components in a single pixel. Instead, the phasor-separation method provides a smoother fluorophores separation, reweighting each pixel by the corresponding fractional components (Fig. 4a).

**Fig. 4.**
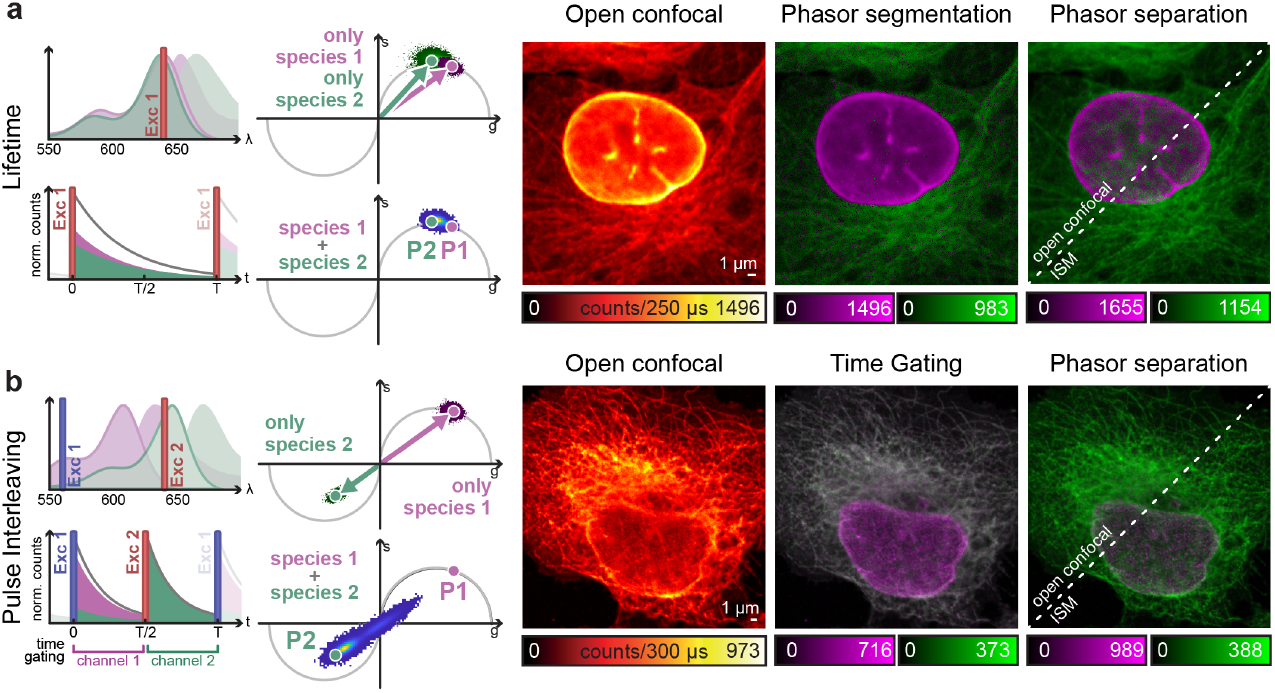
Time-resolved acquisition for multi-colour ISM imaging. (a) ISM imaging of different species, reported by two dyes with similar *spectra* but different fluorescence lifetimes. (b) Pulse-interleaving ISM imaging of different species, reported by two spectrally different dyes. Left: schematic diagrams of (top) the absorption/emission *spectra* of fluorophore 1 (magenta) and 2 (green), and (bottom) of the detected signal over time. The excitation lines are also represented. Right: phasor representation of (top) two measurements obtained with only one species labelled at a time, and (bottom) of the measurement with both species labelled. Left image: raw open confocal image. Central image: open confocal result obtained by (a) segmenting the phasor space in two regions, related to the first and the second fluorophore, or (b) by temporal gating of the detected signal in two temporal channels, related to the first and the second excitation event. Right image: open confocal and ISM results obtained by phasor separation.

An alternative approach for simultaneous multi-species imaging with a single SPAD array detector is the pulsed interleaved excitation (PIE) technique. Thanks to this latter, we can separate the signal from multiple probes based on their differences in the absorption spectra. The PIE technique is particularly convenient if the probes have different absorption spectra but similar emission spectra. It consists of a sequential excitation at different wavelengths, synchronized with the signal recording. This strategy reduces the spectral emission cross-talk without significantly reducing the acquisition speed since the excitation repetition rate can be as fast as tens of MHz. We achieve PIE by synchronizing several pulsed lasers with different colours. The pulses from the different lasers are alternated with a delay much longer than the probe’s fluorescence lifetime to ensure that a dye emits all the photons before exciting the other dye. We record the photons’ arrival time information with respect to the excitation pulse, enabling the generation of different images obtained by binning the photons in different temporal windows – one for each species [33]. This method – known as time-gating – fails if the absorption spectra of the probes overlap since one fluorophore could be excited by multiple excitation colours. Harnessing the phasor information, we remove the cross-talk by unmixing the contributions of different fluorophores by decoding the absorption information encoded in the fluorescence dynamics. To demonstrate the feasibility of the proposed approach, we performed PIE imaging of a fixed HeLa cell (Tubulin labelled with ATTO 647N and nuclear-pore complexes with Abberior STAR 580). The dyes ATTO 647N and STAR 580 have a substantial spectral absorption overlap. Thus, a time-gating approach alone cannot effectively separate the structures labeled with the two dyes. In particular, ATTO 647 can be excited by the lasers at 560 nm and 640 nm, contributing to the counts of the window assigned to STAR 580. To calculate the phasor components of each species, we first imaged two samples with individual labeling. Then, we calculated the phasors along the complete excitation period (*∼*20 MHz). Thus, the photon-arrival histogram contains the fluorescence excited by the green and red pulses. The phasor of STAR 580 (excited by the laser at 560 nm) is localized in the universal semicircle of the first quadrant of the complex plane. Indeed, STAR 580 cannot be excited by red light and emits fluorescence with single exponential decay. The dye ATTO 647N is primarily excited by the red laser but also partially excited by the green laser. The corresponding fluorescence decay is the linear composition of two exponential decays separated in time by half of the excitation period. Since the second excitation, caused by the green laser, is more effective than the first one, the phasor is localized in the third quadrant of the complex plane. Thus, the absorption cross-talk is naturally encoded in the phasor plots of the two species. The phasor-separation method can exploit such information to separate the two species, even in the presence of strong cross-talk. For comparison, we also show the results of simple time-gating, in which one of the two channels of the image contains mixed contributions. As expected, time-gating cannot preserve the specificity of the labelling if the dyes have overlapping absorption spectra. Instead, the phasor separation method successfully distinguishes the contributions of each dye (Fig. 4b).

### 2.4 FLISM for Nanoscopy Imaging

Another advanced fluorescence laser-scanning microscopy technique that can significantly benefit from the single-photon imaging paradigm is stimulated-emission depletion (STED) microscopy. In STED microscopy, a second laser beam, the STED beam, induces stimulated emission on the fluorescent probe. The STED beam is engineered in phase and polarization to generate a doughnut-shaped intensity distribution at the focus. By spatially overlapping the foci of the depletion and excitation beam, the probed region of the laser-scanning microscope reduces in size well below the diffraction limit: the higher the STED beam intensity, the smaller the probed region [34]. We have recently demonstrated that combining STED microscopy with ISM allows reducing the intensity of the STED beam to achieve a target resolution [10]. This benefit is maximal at low STED beam intensity and vanishes for high-intensity values. However, live-cell imaging is typically performed at reduced STED beam intensity to mitigate photo-toxicity, thus making STED-ISM attractive for gentle live-cell imaging. Furthermore, the additional spatial information the detector array provides enables effective background removal [10].

In the context of gentle live-cell STED microscopy, a widely used approach to reduce the STED beam intensity is time-resolved STED microscopy [27, 35, 36] – now implemented in all commercial instruments. Time-resolved STED microscopy is a class of implementations that leverages the relation between fluorescence depletion and fluorescence lifetime. Indeed, stimulated emission opens a new relaxation pathway for the probe, whose rate is proportional to the intensity of the STED beam. Thereby, the higher the intensity, the higher the efficiency of depletion, and the shorter the probe’s fluorescence lifetime. Because the STED beam intensity at the focus is shaped as a doughnut, it induces a fluorescence lifetime spatial signature: the fluorescence lifetime is the shortest at the periphery and unperturbed at the centre of the probed region, where the intensity of the depletion beam is at its maximum and minimum, respectively. Time-resolved STED microscopy harnesses the spatial dependency of fluorescence lifetime to distinguish the fluorescence signal generated from the probed region’s centre or periphery. The result is a smaller effective point spread function (PSF). Thus, the resolution is enhanced without increasing the STED beam intensity.

Time-resolved STED microscopy initially used a time-gated detection [35, 36] to remove the short-lived fluorescent signal. The fluorescence signal is recorded only after a fixed delay (a fraction of the probe’s natural fluorescence lifetime) from the excitation events triggered by the pulsed laser. The longer the delay, the smaller the effective probed region. However, time-gated detection also rejects part of the fluorescence signal generated from the centre, thus reducing the SNR. Given the photon-arrival time histogram of the probed region, a computational alternative can solve the hardware time-gated detection limitations, providing the same resolution improvement but without a strong SNR reduction. In particular, the SPLIT method can separate the long-lived and short-lived fluorescence signal generated from the inner and outer parts of the STED microscopy probed region, respectively [25, 26, 37, 38]. The SPLIT method represents the two decays as a linear combination of phasors. Thus, the unmixing problem of the different components is solved by the SPLIT method by inverting a simple linear system (see Supplementary Note C and Suppl. Fig. D6-D7).

Here, we used our proposed DFD-DAQ module to introduce time-resolved STED-ISM. In particular, we combined APR-based STED-ISM with the SPLIT approach. We call this combination SPLIT-STED-ISM. We first applied the SPLIT approach on the 5D raw STED photon-counts map *i*(**x**_*s*_, **x**_*d*_, *t*), to obtain a set of images *i*(**x**_*s*_, **x**_*d*_) with increased resolution. Then, we used the APR method to achieve the final SPLIT-STED-ISM result, with further increased resolution and SNR with respect to the original raw STED measurement. The APR algorithm calculates the shift vectors directly from the data, thus adapting toward the power of the STED beam. Notably, the shift vectors retrieved before and after applying the SPLIT algorithm differ due to the effective separation of the early and late photons (Supp. Fig. D9). Thus, resolution and SNR are maximized if APR is applied after SPLIT.

We tested the new SPLIT-STED-ISM method on fixed HeLa cells with ATTO 647N labeled tubulin. The longer the duration of the STED pulse, the more useful SPLIT is in removing the incomplete depletion background at the periphery of the probed region [27]. In a practical scenario like ours, the pulse width is non-negligible with respect to a probe’s fluorescent lifetime – about 600 ps to a few nanoseconds. Thus, combining STED-ISM with SPLIT produces a high-contrast and high-quality image (Fig. 5a). Finally, we complemented nanoscopy with multi-species imaging. In detail, we used the PIE approach combined with STED microscopy. We separated the different fluorophores (STAR RED for nuclear pore complexes and STAR ORANGE for the Golgi apparatus) through time-gating – given the lack of fluorophores cross-talk – and applied the SPLIT algorithm to each channel to improve both resolution and SNR. The result demonstrates the vast capabilities of our platform (Fig. 5b).

**Fig. 5.**
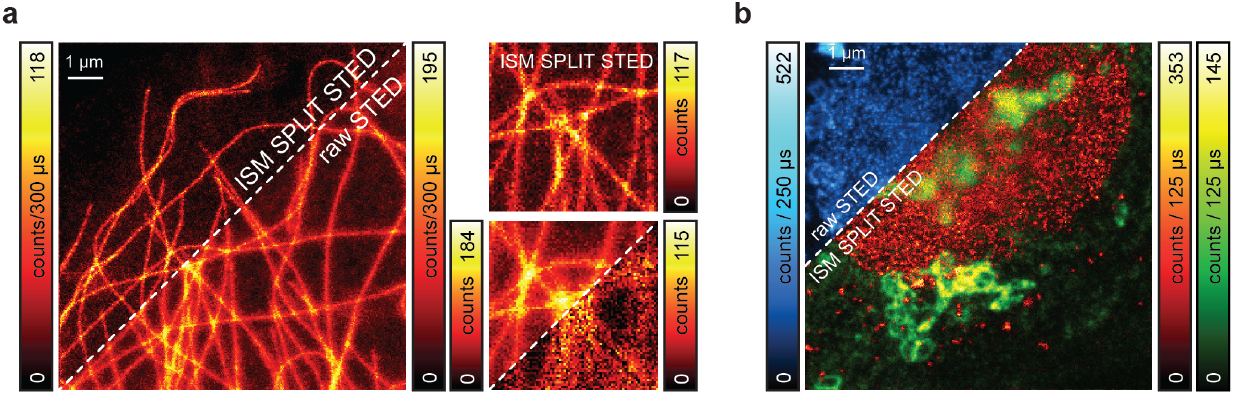
SPLIT-STED-ISM. (a) Side-by-side comparison of raw STED imaing and SPLIT-STED-ISM imaging of the cytoskeleton network. The insets show magnified details of the network. Also the incomplete depletion background is visualised (bottom inset, right corner). (b) multi-species SPLIT-STED-ISM imaging. Pulse-interleaving excitation and time-gating detection allows for reconstructing two datasets – one for each fluorophores – on which performing SPLIT. The upper corner shows the raw STED image. The lower corner the final multi-color SPLIT-STED-ISM image (green: Golgi apparatus, red: nuclear pore complex protein).

## 3 Methods

### 3.1 Microscope Architecture

For this work, we updated the STED-ISM setup described previously [10], adding the possibility to excite the sample also with a green laser beam and to record the photon-arrival time histograms (Suppl. Fig. D1). Briefly, the excitation beams were provided by two triggerable pulsed (*∼* 80 ps pulse-width) diode lasers (LDH-D-C-640 and LDH-D-C-560, Picoquant) emitting at 640 nm and 560 nm, respectively. The STED beam was provided by a triggerable sub-nanosecond (*∼* 600 ps pulse width) pulsed fibre laser (Katana 775, OneFive) emitting at 775 nm. We adjusted the laser power for all beams using their respective drivers and control software. For the 640 nm and 516 nn laser beams, we can further modulate and adjust the power by using acoustic optical modulators (AOM, MT80-A1-VIS, AAopto-electronic), which also act as fast shutters. All laser beams are coupled into a different polarising-maintain fibre (PMF) to transport the beams to the microscope easily. In all cases, we used a half-wave plate (HWP) to adjust the beam polarization parallel to the fast axis of the PMF. For the STED beam, we used a PMF designed explicitly for high-power applications (QPMJ-A3HPM,3S-633-4/125-3-7-1, AMS Technology). The STED beam emerging from the PMF is collimated, filtered in polarization by a rotating Glan–Thompson polarizing prism and phase-engineered through a polymeric mask imprinting 0–2*π* helical phase-ramps (VPP-1a, RPC Photonics). We rotated a quarter-wave plate and a half-wave plate to obtain circular polarization of the STED beam at the back aperture of the objective lens. A set of dichroic mirrors allows the combination of all laser beams (i.e., excitation and STED) and the separation of the fluorescence signals. Two galvanometer scanning mirrors (6215HM40B, CT Cambridge Technology), a scan lens and a tube lens – of a commercial confocal microscope (C2, Nikon) – deflect and direct all the beam towards the objective lens (CFI Plan Apo VC 60×, 1.4 NA, Oil, Nikon) to perform the raster scan on the specimen. The fluorescence light was collected by the same objective lens, descanned, and sent to the detector. A 300 mm aspheric lens (Thorlabs) focuses the fluorescence light into a completely open pinhole plane generating a conjugated image plane with a magnification of 300× . A telescope system, built using two aspheric lenses of 100 mm and 150 mm focal length (Thorlabs), conjugates the SPAD array with the pinhole and provides an extra magnification factor. The final magnification on the SPAD array plane is 450×, thus the size of the SPAD array projected on the specimen is ∼1.4 A.U. (at the far-red emission wavelength, i.e. 650 nm). Two notch filters (561 nm and 640 nm) and a short pass filter (720 nm) reject the light from the laser beams reaching the image plane (i.e., before reaching the SPAD array detector). Depending on the experiments, fluorescence light can be selected by using a bandpass filter (590/50 nm and/or 685/70 nm). The setup mounts a 5 × 5 asynchronous read-out SPAD array detector (PRISM-light kit, TTL version, Genoa Instruments). Every photon detected by any of the 25 elements of the SPAD array generates a TTL signal that is delivered through the dedicated channel (one channel for each sensitive element of the detector) to a multifunction FPGA-based I/O device (NI USB-7856R from National Instruments). The board acts both as DFD-DAQ system and control unit. A LabVIEW-based software, inspired by the Carma software [39], implements the DFD and microscope control modules. Specifically, the software controls the entire microscope devices needed during the image acquisition, such as the galvanometric mirrors, the axial piezo stage, and the acousto-optic modulators (AOMs), builds the 5D photon-counts map *i*(**x**_*s*_, **x**_*d*_, *t*), and visualises the images (e.g., the image obtained from the integration along the *x*_*S*_ and *t* coordinates). Furthermore, the control unit provides the three synchronised signals for triggering the excitation and the STED beams (i.e., *f*_*exc*_ = 2*f*_560_ = 2*f*_640_ = *f*_775_). Each synchronisation signal is sent to a different picosecond delayer to implement the pulse sequence for pulse-interleaving excitation and for an efficient stimulated depletion process. When necessary, the delayer also converts the TTL from the control unit to a NIM signal for the laser driver.

### 3.2 Sample Preparation

#### 3.2.1 Quenched fluorescent solutions

For spectroscopy measurements (Fig. 2 c), we prepared different solutions of fluorescein (46955, free acid, Sigma-Aldrich, Steinheim) at different concentrations of the quencher salt potassium iodide (60399-100G-F, BioUltra, ≥ 99.5% (AT), Sigma-Aldrich). We first dissolved fluorescein from powder into DMSO (Sigma-Aldrich), and we further diluted it to a 1:1000 v/concentration by adding ultrapure water. Then, we diluted the solution at different volume ratios with the potassium iodide quencher (1:2, 1:4, 1:8, 1:16, 1:32, 1:64). All samples were made at room temperature. A new sample solution was prepared before each measurement. Nevertheless not optimal; we excited the sample using the 560 nm laser beam.

#### 3.2.2 Fluorescent beads

To qualitatively characterise the spatial resolution enhancement of ISM and STED microscopy (Suppl. Figs. D3, D6) and we used a commercial sample of ATTO647N fluorescent beads with a diameter of 23 nm (Gatta-BeadsR, GattaQuant).

#### 3.2.3 Fixed cell imaging

To validate our DFD-DAQ system for FLISM imaging (Fig. 3 a,b, Suppl. Fig. D4), multi-species imaging based on pulse-interleaving excitation (Fig. 4 b), and single-colour STED imaging (Fig. 5 a), we used fixed Hela cell labelled to visualise *α*-tubulin and Nup-153 (nuclear pore complexes). HeLa cells were seeded on coverslips in a 12-well plate (Corning Inc., Corning, NY) and cultured in Dulbecco’s Modified Eagle Medium (DMEM, Gibco^™^, ThermoFisher Scientific) supplemented with 10% fetal bovine serum (Sigma-Aldrich, Steinheim, Germany) and 1% penicillin/streptomycin (Sigma-Aldrich) at 37 °C in 5% CO_2_. After 24 hours, cells were incubated in a solution of 0.3% Triton X-100 (Sigma-Aldrich) and 0.1% glutaraldehyde (Sigma-Aldrich) in BRB80 buffer (80 mM Pipes, 1 mM EGTA, 4 mM MgCl_2_, pH 6.8, Sigma-Aldrich) for 1 min. After fixation with a solution of 4% paraformaldehyde (Sigma-Aldrich) and 4% sucrose (Sigma-Aldrich) in the BRB80 buffer for 10 min, cells were washed three times for 15 min in phosphate-buffered saline (PBS, Gibco^™^, ThermoFisher Scientific). Cells were incubated in a 0.25% Triton-X-100 solution in BRB80 buffer for 10 min, then washed three times for 15 min in PBS. After 1-hour incubation in blocking buffer (3% bovine serum albumin (BSA, Sigma-Alrich) in BRB80 buffer), fixed HeLa cells were incubated with monoclonal mouse anti-*α*-tubulin (1:1000, Sigma-Aldrich) and rabbit anti-Nup153 (1:500, ab171074, Abcam, Cambridge, UK) antibodies diluted in the blocking buffer for 1 hour at room temperature. Anti-*α*-tubulin and anti-Nup153 antibodies were respectively revealed by ATTO647N goat anti-mouse IgG (1:400, Sigma-Aldrich) and STAR580 goat anti-rabbit IgG (1:400, Abberior GmbH, Göttingen, Germany) diluted in BRB80 buffer. After 1 hour of incubation, cells were rinsed three times in PBS for 15 min. Finally, coverslips were mounted onto microscope slides (Avantor, VWR International, Milano, IT) with 2,2-thiodiethanol (Sigma-Aldrich).

To validate our DFD-DAQ system for multi-species imaging based on fluorescence lifetime (fig. 4 a), we used fixed MCF10A labelled to visualise *α*-Tubulin and Lamin A. MCF10A cells plated on coverslips coated with the 0,5% w/v pork gelatin (G2500, Sigma-Aldrich) were washed 3x with pre-warmed PBS, fixed with formaldehyde solution (3,7% methanol free) for 15 min, and washed 3x with PBS. After the subsequent blocking with the blocking buffer composed of 0,1% Triton X-100 and 3% w/v of bovine serum albumin (BSA) for 1h, samples were incubated with the following primary antibodies for 1h: mouse anti-*α*-Tubulin (1:1000, T5168, Sigma-Aldrich) and rabbit anti-Lamin A (1:1000, ab26300, Abcam). The samples were washed 2x with the blocking buffer and incubated for 45min in the dark with the following secondary antibodies: goat anti-mouse STAR RED (1:200, Abberior GmbH, Göttingen, Germany) and anti-rabbit STAR635 (1:200, Abberior GmbH, Göttingen, Germany). After washing the samples 5x with PBS, they were mounted with the TDE mounting medium. Additionally, single staining experiments were performed in order to establish the fluorescence lifetime in the cell structure of interest (tubulin or lamin A) separately. All the steps were performed at room temperature, unless otherwise stated.

To validate our DF-DAQ system for multi-colour STED-ISM imaging (fig. 5 b), we used a ready-to-image sample kit (Imaging set for STED@775 nm, Abberior). The microscope slide contains fixed mammalian cells immunostained for a nuclear pore protein with STAR RED (Abberior GmbH, Göttingen, Germany) and the Golgi apparatus protein giantin with STAR ORANGE (Abberior GmbH, Göttingen, Germany).

#### 3.2.4 Live cell imaging

For FLISM functional live-cell imaging, we used Hela cells (Fig. 3c). HeLa cells were seeded on *μ*-Slide 8 well (Ibidi GmbH, Gräfelfing, Germany) and cultured in Dulbecco’s Modified Eagle Medium (DMEM, Gibco^™^, ThermoFisher Scientific) supplemented with 10% fetal bovine serum (Sigma-Aldrich) and 1% penicillin/streptomycin (Sigma-Aldrich) at 37°C in 5% CO_2_. After 24 hours, the growth medium was removed and cells were incubated with 1 μM CellBrite^™^ NIR680 (Biotium, Fremont, CA, USA) for 20 min. After removing the staining medium, HeLa cells were rinsed three times with fresh growth medium; cells were incubated at 37 °C for 5 min between each rinse. Finally, HeLa cells were imaged in Live-Cell Imaging Solution (ThermoFisher Scientific).

## 4 Discussion

We presented an FPGA-based multi-channel DAQ system tailored for fluorescence laser-scanning microscopy with asynchronous read-out SPAD array detectors. This system uses the DFD principle to implement 25 low-resource TDCs. Using limited resources of the FPGA, our architecture enables the integration of the DAQ-DFD module and the microscope control unit into the same FPGA-based board, greatly reducing the microscope cost and complexity. These benefits come at the cost of a lower temporal precision and sampling (*∼* 2 ns and *∼* 400 ps, respectively) than other FPGA-based TDC architectures, which recently has been shown to reach ten picoseconds precision [40]. Nonetheless, the specifications of our architecture are fit to implement FLISM, making its numerous application easily accessible. On the other side, a limitation of the current DAQ module implementation is the poor maximum data-transfer rate of the USB 2.0 protocol, which constrains the pixel dwell time of the scanning microscope to *∼* 100 μs. This problem can be solved using a board with similar FPGA but faster data transfer protocols (e.g., USB 3.0 or PCI Express). This board would also open to more channels (e.g., 49 channels) since the bottleneck is not the FPGA resource but the limited data transfer rate.

We used the proposed DFD-DAQ system to implement the well-established FLISM technique and demonstrate novel advanced imaging techniques based on the single-photon laser scanning microscopy paradigm. In particular, we demonstrated a method capable of distinguishing fluorophores on the base of their absorption spectra. We used the pulse-interleaving excitation scheme to encode the spectral information of the fluorophores into the time domain. Successively, we used a phasor-based approach to decode the fluorescence light emitted by the two species. The proposed method automatically corrects the fluorophore excitation cross-talk. Notably, the same approach can also distinguish fluorophores with different lifetimes, setting the basis for multi-species ISM imaging with a single detector. Thus, we effectively increased the portfolio of usable dyes.

We also demonstrated the combination of ISM with time-resolved STED microscopy. In a recent paper, we showed the benefits of combining STED microscopy with the APR method to reduce the STED beam intensity, thus minimising the risk of inducing photodamage [10]. Here, we demonstrate that leveraging single-photon temporal information can further boost the benefits. We proved that the SPLIT method increases the resolution and contrast of the image. Since the algorithm can be applied to each pixel and channel independently, it leaves the dimensionality of the ISM dataset intact. Thus, the processed data are still compatible with the recent and advanced image reconstruction algorithm developed for ISM – such as focus-ISM [10] or multi-image deconvolution [41]. We envision that maximum likelihood reconstruction methods that consider both the spatial and temporal information will emerge as a superior reconstruction tool for time-resolved STED-ISM datasets. Indeed, the spatiotemporal information is encoded in the temporal PSFs: the stimulated emission process introduces a temporal evolution on the effective PSF. Namely, the strongest the effect of the STED beam, the narrower the PSF associated with each scanned image (Suppl. Fig. D8 and D9).

In conclusion, we believe the proposed architecture will make photon-resolved image scanning microscopy easily accessible, paving the way for gentle, versatile imaging at high spatial resolution and information content.

## Declarations

### Supplementary information

A supplementary file accompanies this article. The supplementary information contains theoretical details, additional methods, and additional figures.

## Acknowledgments

The authors thank Dr. Mario Faretta (Department of Experimental Oncology, European Institute of Oncology, Milan, Italy) for kindly sharing the MCF10A cell line; Andrea Bucci (Istituto Italiano di Tecnologia, Genoa, Italy) for the fruitful discussion about the digital frequency domain principle; all members of the Molecular Microscopy and Spectroscopy labs for the many helpful suggestions.

## Authors’ contributions

G.T., M.C., and G.V. conceived the idea. G.T., A.Z., S.P, M.C., and G.V. designed the study. G.T., A.Z., M.D., and M.C. expanded the theory of the digital-frequency domain (DFD) and its application in the fluorescence lifetime context. G.T. and M.C. implemented the DFD module for the data-acquisition system. G.T., M.D., and M.C. developed the microscope control unit. G.T. and A.Z. built the image scanning microscope. S.Z. and A.P.M. prepared the biological samples. G.T. performed the experiments. G.T. and A.Z. developed the image analysis software. G.T., A.Z., and G.V. analyzed the data with the support of all other authors. G.T., A.Z., and G.V. wrote the manuscript. All authors discussed the results and commented on the manuscript.

## Data availability

The experimental data generated and analysed in this study will be deposited in a publicly available Zenodo database.

## Code availability

The image analysis code used for the current study is available upon request.

## Appendix A DFD - Theory and implementation

### A.1 Working principle of heterodyne acquisition

The core idea of the DFD acquisition is to sample a periodic event at a slightly different periodicity, accumulating signal across multiple periods. Such heterodyne measurement results in the cross-correlation between the sampling window and the event, sampled at a higher rate than it would have been possible with conventional sampling. Indeed, by properly tuning the measurement parameters, the effective time step can be much shorter than the actual duration of the sampling window.

We now analyze the DFD working principle in detail. We excite the fluorescence emission by exciting the fluorophores at the excitation frequency *f*_*exc*_. Then, we impose the sampling frequency *f*_*s*_ to be slightly detuned from the excitation. The cross-correlation frequency *f*_*c*_ is the beating frequency between the sampling and excitation frequency:

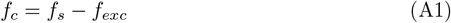

where we assumed *f*_*s*_ *> f*_*exc*_. Then, we impose *f*_*s*_ to be an integer multiple of *f*_*c*_

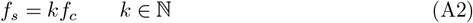

and we define a windowing frequency *f*_*w*_ to be a multiple integer of *f*_*s*_

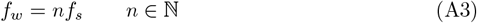

Using the equations above, we find the relation of *f*_*exc*_ with *f*_*w*_ and *f*_*s*_

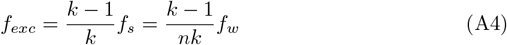

Therefore, the cross-correlation period contains an integer number of excitation and sampling periods

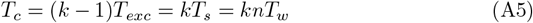

In other words, the excitation and sampling signal are in phase every *T*_*c*_, which is the periodicity of the DFD measurement.

To calculate the arrival time of the detected photons, we define the following counters

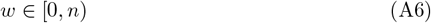

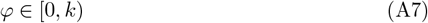

The counter *w* indicates the window of the arrival time, inside a sampling period. The counter *φ* indicates the phase delay between the sampling and excitation signal. Thus, the sampling time with respect to the excitation is

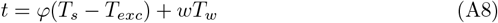

The first term represents the delay between excitation and sampling periods (the time interval between the dashed line and the following exponential decay, in Fig. 1b). The second term represents a time advancement, given by the position of the sampling windows with respect to the sampling clock (the time interval between the dashed line and the window with index *w*, in Fig. 1b). It is useful to rewrite the above equation using a single period as the unit of time

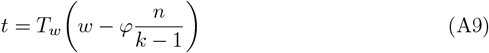

Note that we are sampling a signal with period 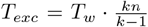. Thus, we take the remainder of the division of the time delay to the period of the signal. Normalizing the result by *T*_*w*_, we get the following dimensionless index

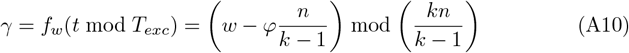

it is useful to multiply both operands by (*k −* 1) to get an integer index

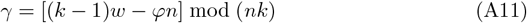

which ranges from 0 to *nk −* 1. The corresponding photon arrival time is

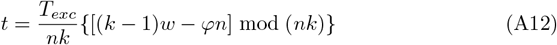

The above equation enables us to sample the excitation period with a step of 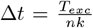. However, in a practical scenario, it could be desirable to trade temporal precision for additional photon counts per time bin. To achieve this goal, we require the windowing period to be an integer multiple of the delay between excitation and sampling periods

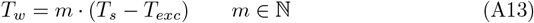

The above condition implies

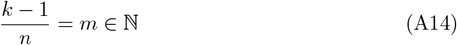

If the above condition applies, then equation A11 can be rewritten as

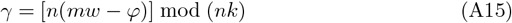

Notably, the time indices can now only be multiples of *n*. In other words, the sampling step is now 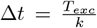. Thus, we further simplify the time index by dividing both operands by *n*

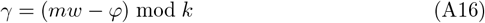

Each index calculated with the above equation is repeated *n* times. Thus, we traded the sampling step size of a factor of *n* to gain the same factor in terms of photon counts. The obtained redundancy comes from the fact that now that the full sampling is repeated *n* times in the *T*_*c*_ period. In other words, the DFD period is now decreased to *T*_*c*_/*n*.

Assuming that *k ≫* 1, we finally obtain

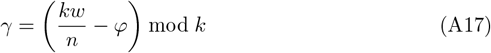

which takes integer values from 0 to *k −* 1. The time index can be reconverted into a time value, knowing that the time interval spans an excitation period. Namely, the photon arrival time is

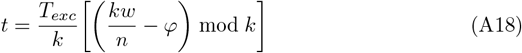

Shifting the photon arrival time index by one window length might be desirable for visualization purposes. Thus, equation A11 becomes

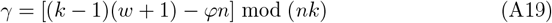

### A.2 Time-Resolved Multi-Channel Data Acquisition System

We upgraded our previously reported acquisition and controlling platform [6] with a multi-channel Digital Frequency Domain (DFD) architecture. The platform is built by two main units: (i) a real-time data-acquisition and controlling module written in LabView FPGA, implemented on a commercially available FPGA board (NI USB-7856R from National Instruments), and (ii) a module running on the host computer, for the user interface and data input-output.

The main tasks of the real-time FPGA platform are to control all the components of the microscope (i.e., lasers, galvanometric mirrors and piezo-electric stage) and to acquire the 25 digital signals from the SPAD array with a temporal precision of up to 400 ps. The platform controls the galvanometric mirrors and the z-axis piezoelectric stage with analog voltage signals. The platform also triggers the excitation and STED pulses through digital signals. Notably, the user can freely define an excitation laser sequence of up to four excitation events per cycle from up to four excitation lasers (in this work, we used up to two excitation lasers and one STED laser). Our design enables a straightforward Pulsed Interleaving Excitation (PIE) implementation. The most significant task of the real-time module is to acquire all the signals from the SPAD array detector with a sub-nanosecond temporal resolution. To this aim, we exploited the DFD principle, leveraging the real-time capabilities of the FPGA technology [23].

For ease of implementation, we chose the ratio *k/n* to be an integer. Therefore, the term *kw/n* in equation A17 can be replaced by a single counter with step *k/n*. More in detail, we used two combinations of parameters, that we refer to as the 20 MHz and the 40 MHz implementations. The chosen parameters can be seen in Table A1, and in

**Table A1.** DFD parameters. From left to right: excitation frequency, sampling frequency, number of sampling periods per cross-correlation cycle, number of windows per sampling period, duration of the sampling window, and the width of the sampling step.

| Implementation | $f_{exc}$ [MHz] | $f_s$ [MHz] | k | n | $T_w$ [ns] | $\Delta t$ [ps] |
| --- | --- | --- | --- | --- | --- | --- |
| 20 MHz | 21 | 21.18 | 120 | 24 | 1.97 | 397 |
| 40 MHz | 39.33 | 40 | 60 | 12 | 2.08 | 424 |

both cases, we obtain a sampling step of Δ*t* = *T*_*exc*_*/k ≈* 400 ps. At the FPGA level, the platform implements a Single-Cycle Timed Loop (SCTL), running at *f*_*SCT L*_ = 12*f*_*exc*_, to generate the TTL signals to trigger up to four lasers. Independently, the data acquisition loop monitors the 25 signals from the SPAD array detector at a frequency *f*_*w*_ = *nf*_*s*_. Each time a photon hits a given sensitive element, the data acquisition loop detects the rising edge of the corresponding TTL signal and increments the correct cross-correlation histogram bin accordingly, accounting the phase relation between the sampling and the excitation frequencies thanks to the phase counter *φ*.

The design choice to generate all the frequencies needed from the same FPGA oscillator implies the compatibility of our platform only with lasers that can be triggered externally. Although this strategy may limit our implementation in its current version, it also results in a significantly simplified controlling architecture, since it does not rely on external signals from the laser sources. Thus, no phase-locking circuits are needed. Additionally, the use of a single oscillator to generate all the frequencies increases the robustness of the platform. However, it does not allow to set the phase relation for the two frequencies at the beginning of the measurement. Indeed, one would like to start every measurement in the same phase condition – e.g., the rising edges of both *f*_*SCT L*_ and *f*_*w*_ being simultaneous – since the value of the phase index *φ* is always set to 0 for the first sampling period. In reality, this condition is not met, and multiple measurements produced at the same experimental conditions are not temporally aligned. To address this problem, we decided to implement an additional virtual data channel. We measure a digital signal provided by the FPGA itself at the frequency *f*_*exc*_, sampled by the same SCTL accounting for the real signals from the SPAD array. Since its temporal position depends solely on the initial phase relation between *f*_*SCT L*_ and *f*_*w*_, it provides a reference for registering different measurements.

At the end of each pixel dwell time, the cross-correlation histograms for each sensitive element of the detector are sent to the host computer *via* the USB 2.0 connection, and are saved by a dedicated LabView module on a binary file. The chosen communication protocol consists of 16 bits per each cross-correlation histogram bin, per each channel. Such protocol enables high-photon flux regimes, but sets a lower limit for the duration of the dwell time to avoid data loss (in typical conditions, 125 μs). It is important to highlight that this limitation is purely technical, and mostly related to the communication bus equipped with the FPGA card. Indeed, implementing the firmware on more recent FPGA platforms (e.g., equipped with Thunderbolt 3 cards) would result in a performance boost up to 80× in data transfer, *de facto* solving the dwell time limitation. Moreover, the negative exponential nature of the fluorescence decays signals suggests the potential implementation of an adaptive communication protocol, which could use different time indices, shortening the minimum pixel dwell time.

## Appendix B Description of fluorescence dynamics in time and frequency

### B.1 Temporal analysis of lifetime data

Following the previous description of the histogram formation with the DFD approach, the fluorescence lifetime histogram *f* (*t*) can be written as

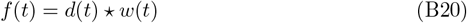

where ⋆ denotes the cross-correlation operator. The fluorescence signal *d*(*t*) is defined as follows

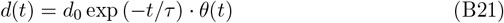

where *θ*(*t*) the Heaviside step function and *τ* the fluorescence lifetime. The sampling window *w*(*t*) is defined as follows

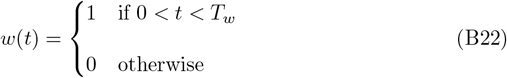

where *T*_*w*_ is the width of the sampling window.

Using the model above, we can analytically compute the cross-correlation

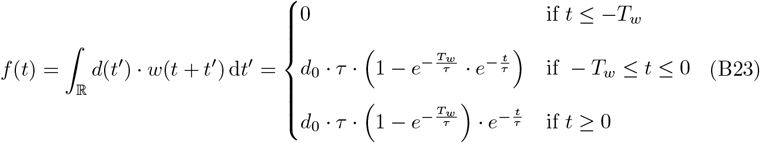

Notably, the result for *t ≥* 0 is still an exponential decay with unaltered lifetime, rescaled by a term that depends on *τ* and *T*_*w*_. The scale factor is

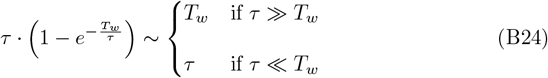

Namely, the contrast of the fluorescence signal linearly deteriorates for decays faster than the sampling window. Instead, the contrast of decays slower than the sampling window is approximately constant.

### B.2 The phasor analysis of lifetime data

Exponential decays are conveniently analyzed in Fourier space. We consider the case of a single exponential

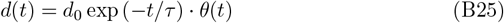

with fluorescence lifetime *τ* . The Fourier transform of the normalized signal is

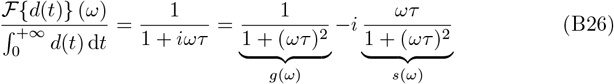

where we defined *g*(*ω*) and *s*(*ω*) respectively as the real and imaginary part of the Fourier transform of the decay. Notably, these quantities are related by the following equation

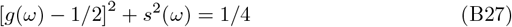

The vector **p** = (*g, s*) is known as phasor and lies on the semicircle of the complex plane described by the above equation, commonly named the *universal circle*. This fact implies that the phasors of single exponential decays are bound to lie on the universal circle. Multi-exponential decays are linear combinations of single exponential decays and their corresponding phasors lie within the universal circle.

From equation B26 it is possible to calculate the estimated lifetime in two ways.

By defining

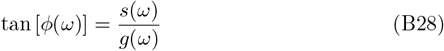

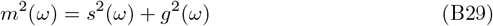

we have that

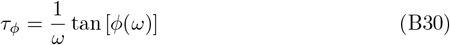

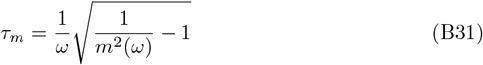

Note that – for single exponential decays – the two estimates of the lifetimes are identical and do not depend on the frequency *ω*.

Sampled data are inherently discrete. Thus, we need to generalize our analysis by writing the phasor coordinates as the real and imaginary part of the discrete Fourier transform (DFT) of the sampled signal:

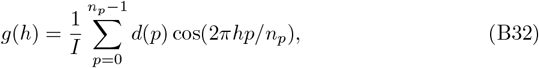

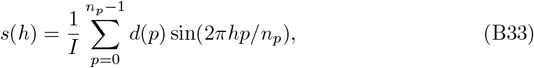

where *I* = Σ_*p*_ *d*(*p*) and *n*_*p*_ is the number of data points. Identifying *ω* = 2*πhf*_exc_, we have the following numerical estimates of the fluorescence lifetime:

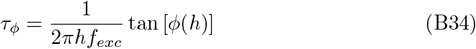

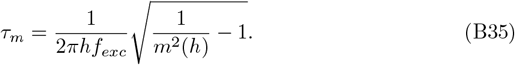

Importantly, phasors can be calculated at any discrete frequency *h*. However, low frequencies carry most of the signal. As such, a typical choice is *h* = 1, commonly referred to as the first harmonic.

All the data analysis presented in this work is performed leveraging a custom-written Matlab application, hosting a graphical user interface to load and manipulate the raw data files generated by the LabView module, taking full advantage of the phasors representation for time-resolved measurements.

### B.3 Calibration of the DFD acquisition platform

In order to retrieve the correct lifetime values from the time-resolved measurements, it is necessary to calibrate the effect of the instrument response on the measurement. Indeed, in a realistic scenario, the impulse response function (IRF) of the microscope is

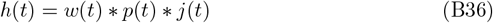

Where *∗* denotes the convolution operator, *w*(*t*) is the DFD sampling window, *p*(*t*) is the envelope of the excitation laser pulse, and *j*(*t*) is the jitter of the detection system. Thus, we can write the fluorescence lifetime histogram *f* (*t*) as

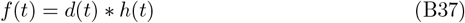

where *d*(*t*) is the fluorescence signal and *h*(*t*) is the IRF of the system. In Fourier space, the same relation can be rewritten using the convolution theorem

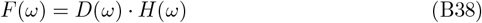

Fixing a single frequency (the one chosen for the phasor calculation) and writing the functions in the exponential form, we have

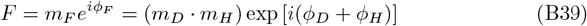

We can estimate the modulus *m*_*H*_ and the phase *ϕ*_*H*_ of the IRF using a sample with a known lifetime, usually a solution of a fluorophore of choice. For such calibration measurement, each of the 25 time-resolved images is integrated along the (*x*_*s*_, *y*_*s*_) coordinates since the temporal fluorescence behaviour and the instrument response are invariant to the scan position. The time-resolved calibration measurement yields to a set [*ϕ*_*C*_(**x**_*d*_), *m*_*C*_(**x**_*d*_)], one phasor for each SPAD element at coordinate **x**_*d*_ = (*x*_*d*_, *y*_*d*_). Similarly, we describe the IRF of each SPAD element as the set [*ϕ*_*H*_ (**x**_*d*_), *m*_*H*_ (**x**_*d*_)] in order to account for potential differences between channels (e.g., different circuitry at the detector level, resulting in different recorded delays for simultaneous photons hitting different channels). Using equations B35 and the known lifetime 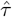, we can estimate the true modulus and phase of the phasor

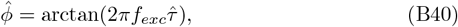

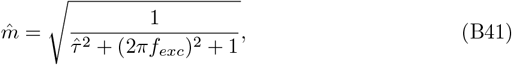

here calculated at the first harmonic, *h* = 1. Thus, we obtain the instrument response as a rotation and a scale in the complex plane.

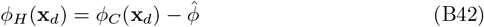

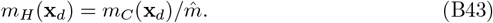

Then, we use the instrument response set to calibrate all the temporal histograms at each position (*x*_*s*_, *y*_*s*_), finally obtaining the correct values [*ϕ*_*D*_(**x**_*d*_), *m*_*D*_(**x**_*d*_)] from the measured values [*ϕ*_*F*_ (**x**_*d*_), *m*_*F*_ (**x**_*d*_)]

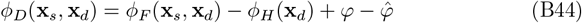

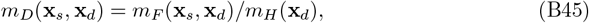

where all the arithmetic operations are intended to be element-wise, and *φ* and 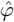 are the phases retrieved from the additional virtual channel, respectively, for the experimental measurement and the calibration measurement. These two terms account for a non-deterministic phase relation between the excitation and the frequency sampling at the beginning of each measurement and cannot be pre-calibrated. In general, the described calibration in the phasor domain allows to retrieve the correct lifetime values for a given measurement – by accounting for the instrument response – and to manipulate the data for a variety of operations directly in the phasor space (e.g., segmentation, decomposition, filtering). Additionally, for this work, it is also beneficial to back-propagate the calibration operation in the image space, since it allows to inherently correct for the temporal differences between different channels of the SPAD array detector. The resulting, “temporally aligned” images may then be fed to analysis pipelines working in the images space (such as the SPLIT-ISM pipeline).

## Appendix C Image processing

### C.1 FLISM data analysis

The acquisition system generates the five-dimensional array *i*(**x**_*s*_, **x**_*d*_, *t*). First, we apply the adaptive pixel reassignment (APR) algorithm [6] to generate the corresponding FLISM image. To do it, we calculate the ISM images by integrating the data over time

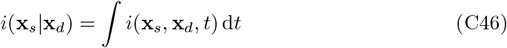

Now, we calculate the phase correlation between the images of the ISM dataset and the one generated by the central element, used as the reference

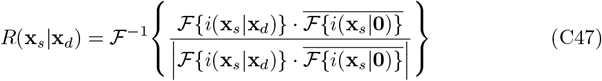

where *F* and *F*^*−*1^ denote the Fourier transform and its inverse, respectively. We measure the shift vectors as the position of maximum correlation

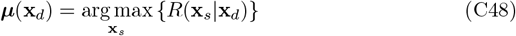

Finally, the FLISM image is calculated by shifting the 5D dataset by the corresponding shift-vectors and by integrating over the detector space

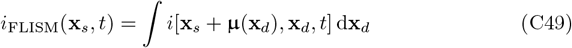

#### C.1.1 Phasor separation

Here, we describe the algorithm we use to separate the contributions of fluorophores with different lifetimes. It consists of a linear method built on the phasor representation of time-resolved fluorescence data. It decomposes the dataset *i*_FLISM_(**x**_*s*_, *t*) pixel-by-pixel by calculating the phasor of the temporal component and unmixing different contributions by inverting a linear model.

For each scan coordinate **x**_*s*_, we assume that the flux of of fluorescence photons contains two components

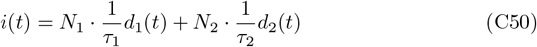

where *N*_1_ and *N*_2_ are the numbers of photons collected with a different lifetime and

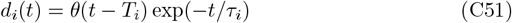

where *T*_2_ = 0 or *T*_2_ = *T*_*exc*_*/*2 for synchronous or pulsed interleaving excitation, respectively. We describe each component using the phasor representation **p** = (*g, s*) and obtain the following linear system

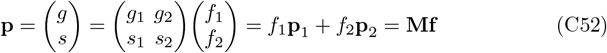

where each phasor **p**_*i*_ represent a single exponential decay and 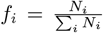 weights that describe the corresponding fractional components. If the matrix **M** is known, it is possible to retrieve the vector of fractional components **f** from a measured phasor **p**.

The linear system described by Eq. C52 solution can be simply solved as follows

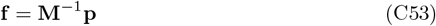

We iterate such procedure for each scan coordinate **x**_*s*_, obtaining the maps of fractional components **f** (**x**_*s*_). Finally, the phasor-separated images are calculated as

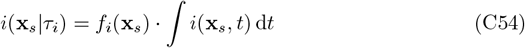

To find the entries of the matrix **M**, we calculated the phasors **p**_1_ = (*g*_1_, *s*_1_) and **p**_2_ = (*g*_2_, *s*_2_) by measuring the fluorescence decay of two calibration samples, each with a single fluorophore. In detail, each phasor has been calculated by averaging the pixels over a scan.

#### C.1.2 Phasor segmentation

To implement phasor-segmentation, we considered in the phasor space the two semi-planes separated by the normal to the vector connecting **p**_1_ and **p**_2_, passing through its middle point. We then generated two images, each containing only the pixels corresponding to phasors in the first or second semi-plane, respectively.

### C.2 SPLIT-STED-ISM data analysis

In order to improve STED-ISM resolution, we used the Separation by Lifetime Tuning (SPLIT) method [25, 26, 42]. Shift-vectors change upon applying the SPLIT algorithm (Suppl. Fig. D6). Thus, we first applied SPLIT to the data for each **x**_*s*_ and **x**_*d*_ coordinate, then applied APR to reconstruct a single image with enhanced resolution.

#### C.2.1 Separation by Lifetime Tuning

The SPLIT algorithm exploits the fluorescence dynamics to separate photons generated from the inner or outer regions of the probed region in STED microscopy. In this case, the flux of fluorescence photons is

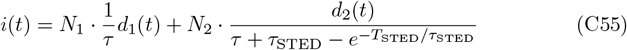

Where

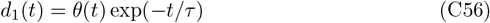

and

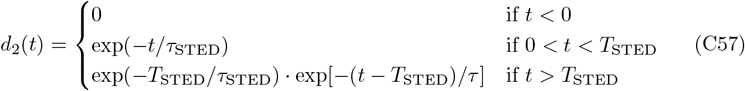

*τ* is the unperturbed lifetime of the fluorophore, *τ*_STED_ is the lifetime shortened by the depletion beam, and *T*_STED_ is the duration of the depletion pulse. Photons emitted from the center of the probed region are unperturbed and decay slower than those originating from the periphery. To distinguish the two families of photons, we used the same procedure described in the previous section. However, in this case, the phasor describing the peripheral photons does not lie on the universal circle but on the trajectory connecting the unperturbed phasor **p**_1_ = (*g*_1_, *s*_1_) to the coordinate (1, 0). In this case, the phasor **p**_1_ = (*g*_1_, *s*_1_) is calculated in the absence of depletion beam, and the phasor **p**_2_ = (*g*_2_, *s*_2_) is manually chosen among the points on the STED trajectory. Thus, we can build the matrix **M** and proceed as described in the previous section.

## Appendix D Supplementary Figures

**Fig. D1.**
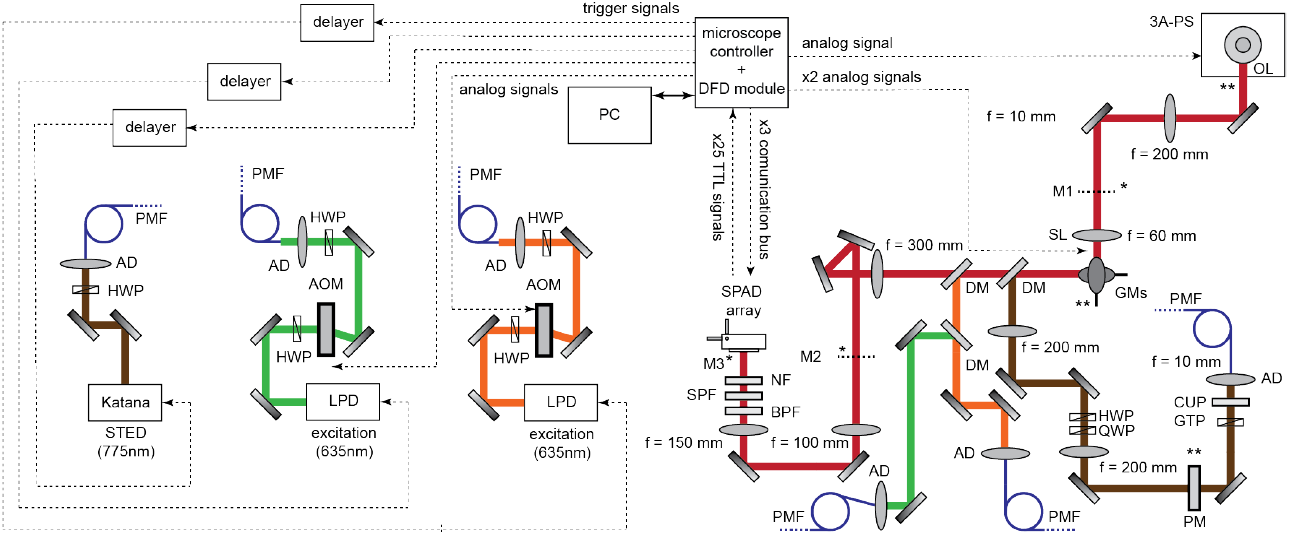
Time-resolved STED-ISM Setup. HWP: half-waveplate; QWP: quarter-wave plate; AOM: acousto-optic modulator; AD:achromatic doublet; PMF: polarized-maintaining fiber; FI: Faraday isolator; 3A-PS: three-axis piezo stage; SL: scanning lens; OL: objective lens; GMs: galvanometric mirrors; DM: dichroic mirror; BPF: band pass filter; CUP: clean-up filter; NF: notch filter; SPF: short-pass filter; APD: avalanche photo-diode; PM: phase-mask; GTP: Glan-Thompson polarizer; . Magnifications: M1 = 60×, M2 = 300×, M3 = 450× . The asterisk denotes the plane conjugate to the image plane. The double asterisks denote the plane conjugate to the objective back-aperture.

**Fig. D2.**
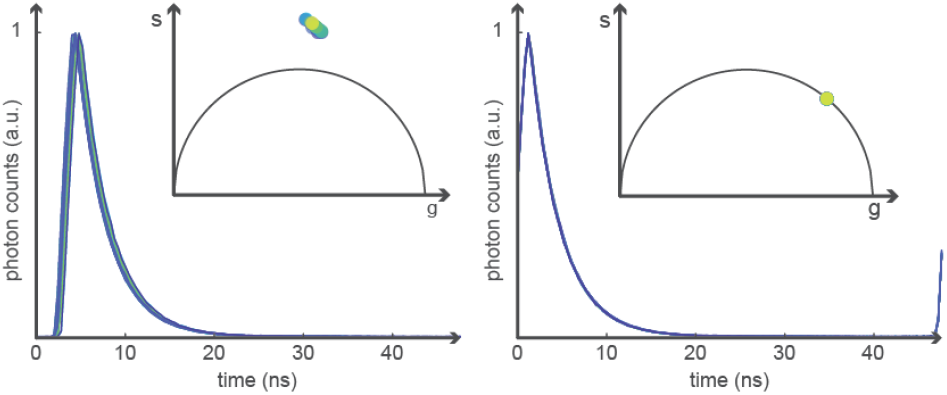
Multi-Channel calibration of the acquisition system. The decays registered by each sensitive element of the SPAD array detector before (left) and after (right) the multi-channel calibration procedure.

**Fig. D3.**
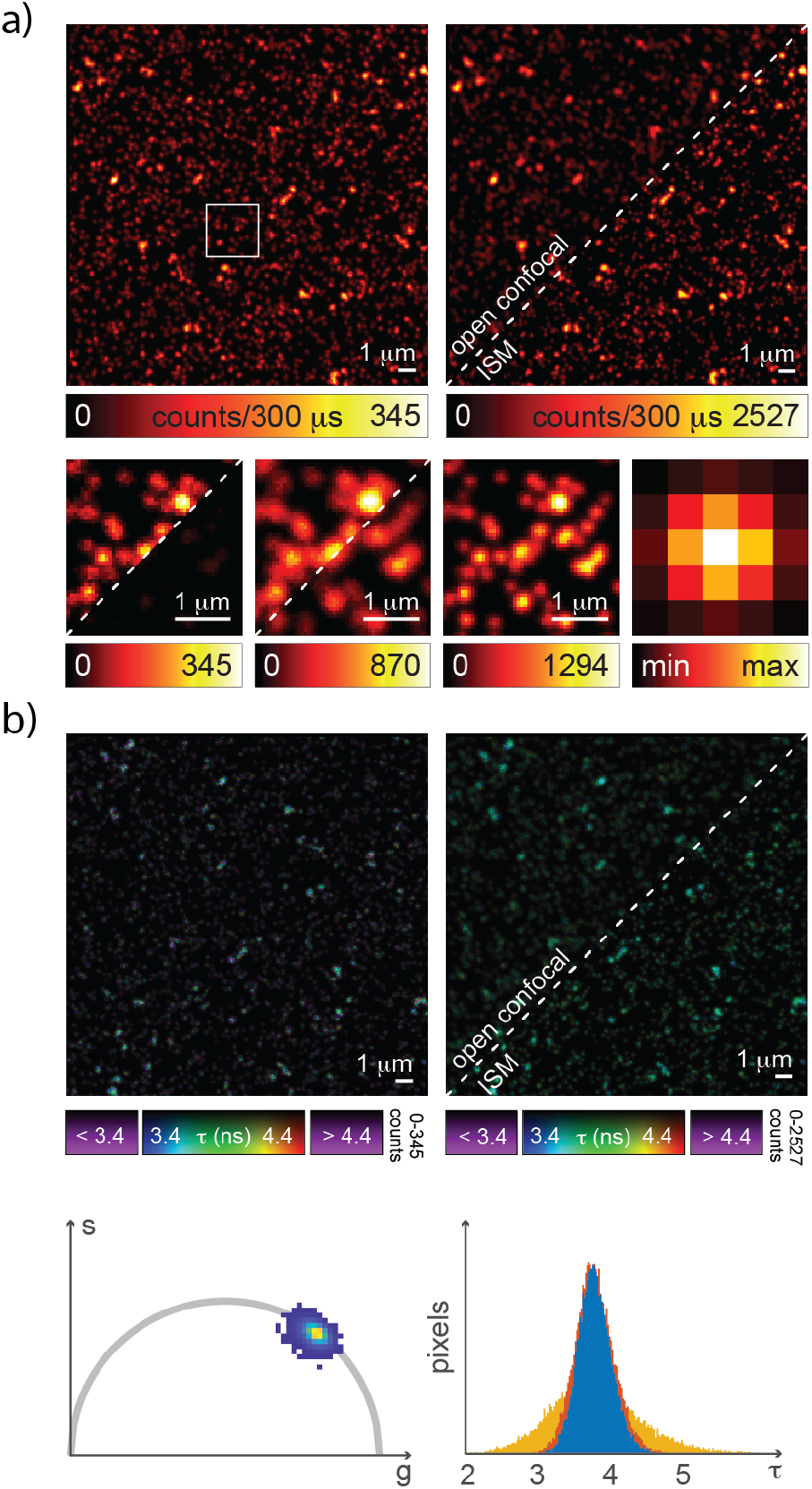
Time-resolved imaging of fluorescent beads. a) Closed confocal (top left, left inset), open confocal (top right, central inset) and ISM imaging (top right, right inset) of fluorescent beads. Right inset: fingerprint matrix. b) Closed confocal (left), open confocal, and ISM lifetime analysis (right) of the same data set in a). Phasor plot for the ISM fluorescence lifetime data set (bottom left) and histograms of the retrieved lifetime values for the different imaging modalities.

**Fig. D4.**
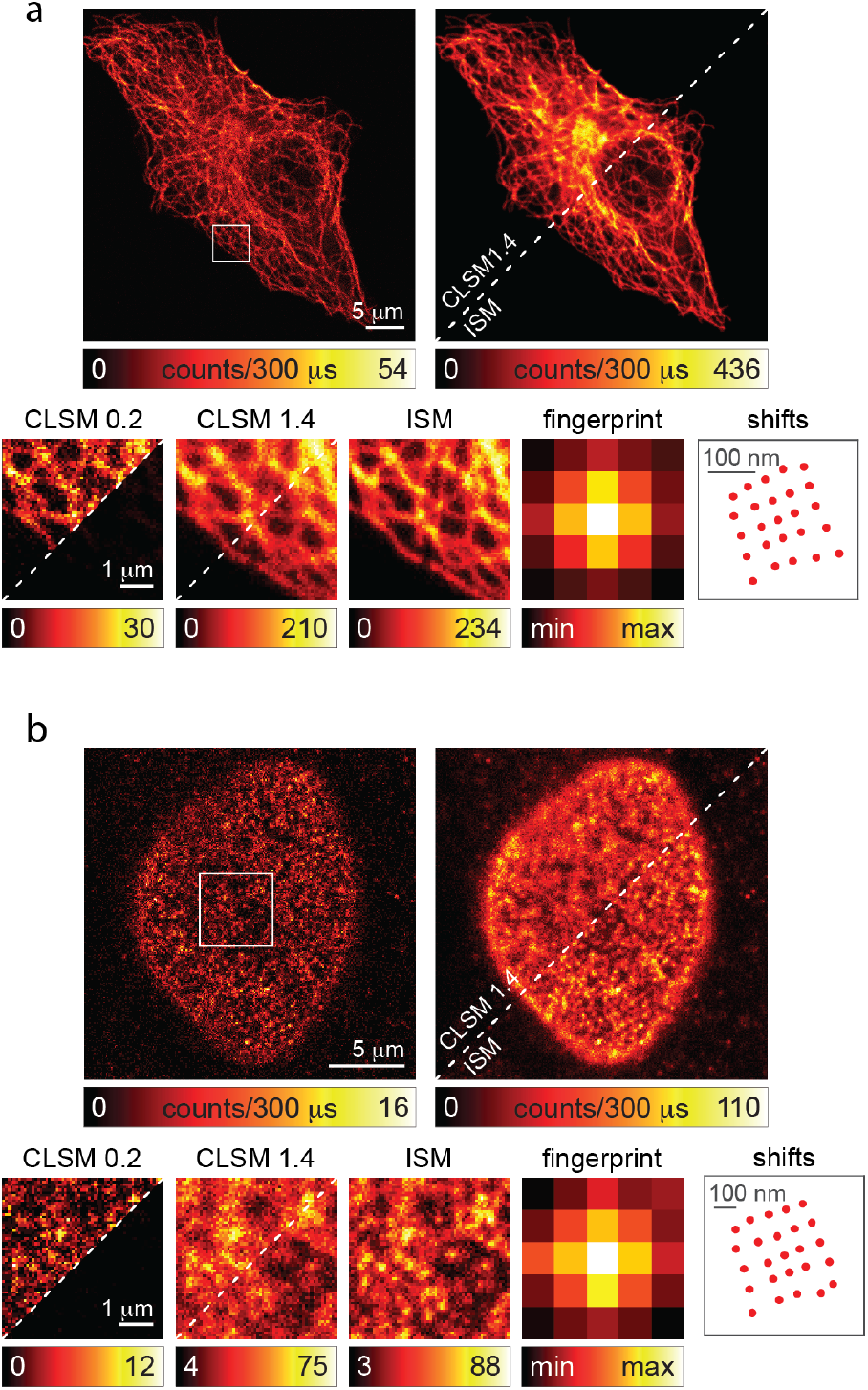
ISM imaging of fixed samples. The panel shows the intensity-based images for the same dataset of Figure 3 (a, b). For imaging of tubulin network (a) and nuclear pore complex (b) the panel shows the close-pihnole image (i.e., central SPAD array element, 0.2. A.U.), the open-pinhole (i.e., the sum of all SPAD array elements, 1.4 A.U.), and the ISM image. The insets shows a detail of the large-field of view image. To highlight the SNR difference between the modality the right corner of the inset are intensity normalised to the ISM counterpart. Fingerprint and shift-vectors are also reported.

**Fig. D5.**
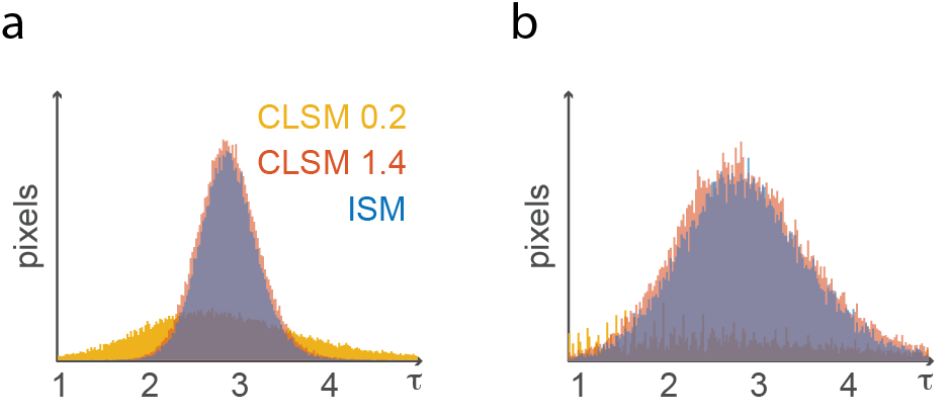
Fluorescence Lifetime histograms for fixed cells. The two histograms represent the fluorescence lifetime values for the tubulin network (a) and the nuclear-pore complex (b) sample. Because of the higher SNR of open-pinhole and ISM images than the close-pinhole images, the relative fluorescence lifetime values show a narrower distribution.

**Fig. D6.**
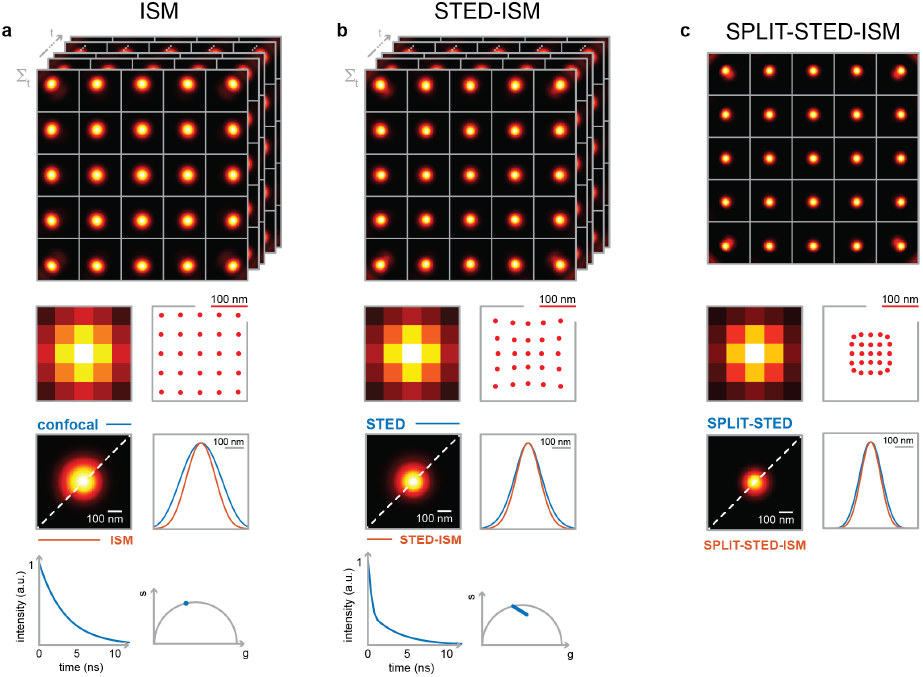
SPLIT-STED-ISM simulations. Simulations of the 5D raw (a) confocal and (b) STED photon-counts map *i*(**x**_*s*_, **x**_*d*_, *t*) recorded by the system when imaging a point source. Top row: representation of the time-dependent PSF for each sensitive element of the detector array. Second row: fingerprint matrix and shift-vectors obtained by integrating each PSF in time. Third row: comparison between (a) raw confocal and ISM or (b) raw STED and STED-ISM. Bottom row: temporal decay of the cumulative PSF, obtained summing all the time-dependent PSFs over **x**_*s*_ and **x**_*d*_ (left); phasor representation of the cumulative PSF obtained summing over **x**_*d*_ only (right). (c) Top: the *i*(**x**_*s*_, **x**_*d*_, *t*) results of the SPLIT approach applied on the raw STED photon-counts map *i*(**x**_*s*_, **x**_*d*_, *t*) in (b). Second row: corresponding fingerprint matrix and retrieved shift-vectors. Third row: comparison with the SPLIT and SPLIT-ISM results – obtained by summing and performing adaptive pixel reassignment, respectively.

**Fig. D7.**
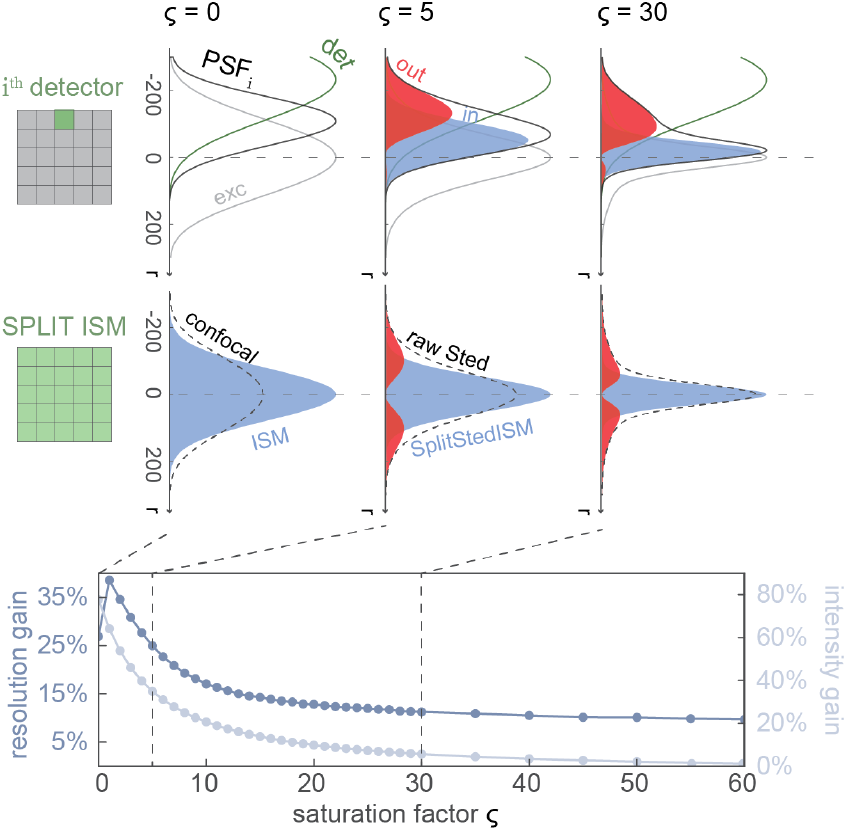
Resolution and intensity gain of SPLIT-STED-ISM imaging. Top row: excitation, detection and effective PSFs of a later detector, displayed for increasing STED saturation factors. In the central and right columns, the high-(blue) and low-(red) resolution contributions are also displayed, as obtained by applying the SPLIT approach. Central row: effects of the adaptive pixel reassignment approach on confocal (left) and STED (center, right) imaging. The ISM (left) or SPLIT-STED-ISM results are displayed in blue. In red, the contribution obtained performing the adaptive pixel reassignment on the low-resolution contributions from all the detectors, as obtained by applying the SPLIT approach. Bottom row: resolution gain – as reported by the Rayleigh value – and intensity gain of the SPLIT-STED-ISM with respect to the raw STED counterpart, with increasing STED saturation factor. The first data point is obtained considering the ISM result and the raw confocal counterpart. The Rayleigh resolution value is obtained using the following strategy: (i) computing the PSF for two point sources at increasing center-to-center distance; (ii) computing the contrast curve as a function of the center-to-center distance. The contrast is defined as (*max − lmin*)*/max*, where *max* is the peak value of the intensity line profiles of the two neighbouring objects, and *lmin* is the value of the local minimum between the two peaks [36]. The Rayleigh resolution is the center-to-center distance for which the contrast reaches the Rayleigh value of 0.26.

**Fig. D8.**
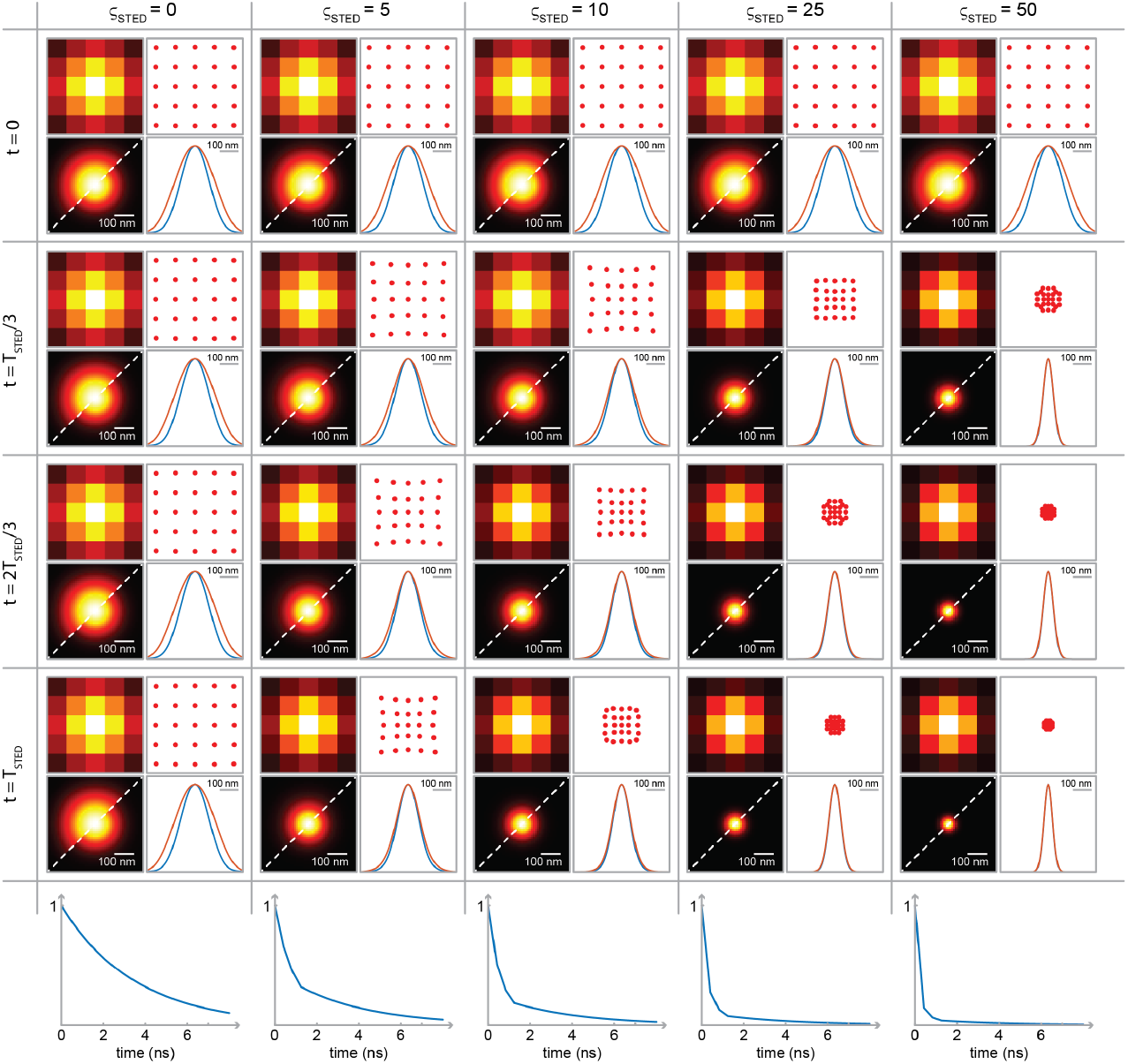
Simulations of time-dependent confocal, raw STED and STED-ISM PSFs. Simulation of the PSFs of a microscope equipped with a detector array (from top to bottom) at different times from the excitation event (*t* = 0) and (from left to right) for increasing STED beam saturation intensities. In each cell: fingerprint matrix describing the distribution of photons on the detector array (top left); shift-vectors retrieved by the adaptive pixel reassignment algorithm (top right); comparison between the open-pinhole (i.e., the sum of all SPAD array elements, 1.4 A.U.) and the STED-ISM PSF (bottom left), and the related line profiles (bottom right). Bottom row: temporal decay of the cumulative PSF for increasing STED saturation factors, obtained by summing all the time-dependent PSFs over **x**_*s*_ and **x**_*d*_. Parameters for the simulation: excitation wavelength: 646 *nm*; depletion wavelength: 775 *nm*; emission wavelength: 669 *nm*; STED pulse duration: 1.25 *ns*; fluorescence lifetime: 3.5*ns*.

**Fig. D9.**
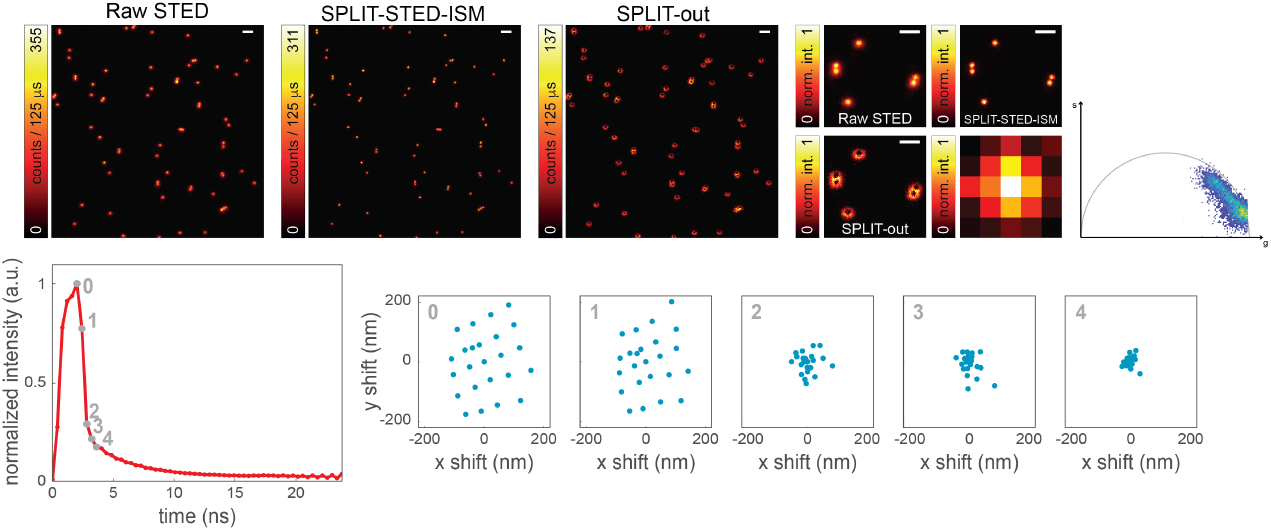
Raw STED and SPLIT-STED-ISM imaging of fluorescent beads. Side-by-side comparison of the Raw STED, the SPLIT-STED-ISM and the SPLIT-out images. The SPLIT approach applied the 5D raw STED photon-counts map *i*(**x**_*s*_, **x**_*d*_, *t*) generates two sets of images *i*(**x**_*s*_, **x**_*d*_): (i) the SPLIT set, on which we used the APR method to achieve the final SPLIT-STED-ISM result; and (ii) the SPLIT-out set, that we integrated over **x**_*d*_ to generate the SPLIT-out image. Right: phasor plot of the raw STED measurement. Bottom, left: temporal decay obtained by integrating the STED photon-counts map *i*(**x**_*s*_, **x**_*d*_, *t*) over **x**_*s*_ and **x**_*d*_. Bottom, right: the shift-vectors for the STED photon-counts map *i*(**x**_*s*_, **x**_*d*_, *t*_*i*_) at increasing delays from the depletion event, as retrieved by the adaptive pixel reassignment algorithm. *t*_0_ = 0 ns, *t*_1_ = 0.4 ns, *t*_2_ = 0.8 ns, *t*_3_ = 1.2 ns, *t*_4_ = 1.6 ns. Image format: 500 x 500 pixels; details format: 100 x 100 pixels; pixel size: 40 nm; excitation power: 3 *μ*W; STED power: 87.5 mW.

